# Vector coordinate transformation in the *Drosophila* navigation system

**DOI:** 10.64898/2025.12.20.695733

**Authors:** Benjamin Gorko, Sung Soo Kim

## Abstract

Animals’ navigational behaviors rely on internal representations of space, yet how these representations guide behaviors is poorly understood. Here, we determined how neurons in the brain of the fruit fly, *Drosophila melanogaster,* transform a world-centric (allocentric) reference frame to one that is body-centric (egocentric), which is essential for navigating to a goal. PFL1 neurons in the fly’s central complex, the navigation center, have two dendrites; one of these receives input from neurons that encode a head direction vector, while the other receives input from neurons in a neuropil called the fan-shaped body (FB). Therefore, we investigated whether PFL1 neurons transform allocentric vectors to an egocentric reference frame. We developed a computational model that shows how PFL1 can transform an allocentric direction vector encoded by FB neurons to an egocentric reference frame by nonlinearly combining it with the head direction input. In tethered flying flies, we simultaneously recorded the head direction input to as well as the output of PFL1, which we used to constrain and fit our model. This allowed us to infer that PFL1 neurons receive a stored direction that persists across behavioral trials for the same fly but appears to be random across flies, fundamental characteristics of a goal. Our results describe how a population of neurons performs a coordinate transformation that can facilitate the pursuit of a goal, a fundamental computation for any neural system containing representations in multiple reference frames.

## Introduction

Successfully navigating in the world reflects the culmination of multiple complex cognitive processes: the brain must perceive its current state, compare it to an internal representation of a goal, and then determine an appropriate next action. Operationally, this requires integrating one’s sense of direction in the world with an internally maintained goal vector to generate new information representing the goal vector relative to the body. Yet the neural circuit mechanisms that transform the reference frame of an internally stored goal vector from allocentric (world-centered) to egocentric (self-centered) are poorly understood.

Neural representations of navigational directions have been identified across species, e.g., head direction cells^1–11^ and goal direction cells^12–17^. In insects, recent physiological and anatomical studies of the central complex (CX, a collection of neuropils at the center of the brain) have revealed how streams of navigational information flow in the CX and how they are integrated and manipulated to generate navigational vectors^18–22^ that include the head direction^2,4^, allocentric locomotion direction^23,24^, goal direction^25–27^, and wind direction^28–31^.

The first ingredient for these vector manipulations in flies is the head direction signal encoded by EPG neurons^2^ (also known as compass neurons) in the ellipsoid body (EB), a donut-shaped CX subregion that comprises a network configuration known as a ring attractor^32–42^. Two copies of the head direction signal are then conveyed to two sides of a handlebar-shaped CX subregion called the protocerebral bridge (PB), where PB-intrinsic interneurons (Δ7 neurons) transform them into two sinusoidal activity profiles^18,23,32^. Nearly all cell types that are postsynaptic to Δ7 and project to the fan-shaped body (FB, another CX subregion) inherit the sinusoidal activity profile. This represents vectors: the amplitude of the sinusoidal profile encodes the vector’s magnitude and the phase of the sinusoidal profile encodes the vector’s direction^18,23,24,29,43,44^. This syntax simplifies the neural implementation of vector computation, similar to the phasor, a mathematical technique often used in physics and engineering^45^.

Intriguingly, some of the FB neurons postsynaptic to Δ7 are involved in generating motor signals for steering while navigating. Specifically, neurons called PFL2 and PFL3 combine information from the PB and FB to transform an immediate goal direction into a steering signal when the fly is walking, playing a critical role in locomotion course correction on short timescales^25,46^.

Here, we report that PFL1 neurons, the remaining uncharacterized class of PFL neurons, transform a vector stored in the FB from an allocentric reference frame to an egocentric reference frame. Although PFL1 neurons have broadly similar morphology and network topology to PFL2 and PFL3, PFL1 neurons are unique in that their output encodes a vector in a two-dimensional Cartesian coordinate system.

We used electron microscopy data to constrain a computational model of the neural circuit structure that underlies PFL1’s input-output relationship, and then we physiologically tested predictions from this model. We found that localized optogenetic stimulation induced steering behaviors, suggesting that the vector stored in the FB represents a flexible, longer-term goal direction and PFL1 neurons convert it into an egocentric motor signal. Thus, PFL1/2/3 neurons collectively convert allocentric vectors to egocentric motor signals on either short-term (PFL2/3) or long-term (PFL1) timescales, depending on behavioral context.

## Results

We analyzed PFL1 neurons’ connectivity using two electron microscopy datasets (the Hemibrain^47^ and the Full Adult Fly Brain (FAFB) dataset^48–50^). PFL1 dendrites tile two neuropils in the fly brain: the protocerebral bridge (PB) and the fan-shaped body (FB) (Fig. 1b). Each PFL1 neuron has a dendrite that occupies a narrow portion of the width of each of these neuropils (Fig. 1b). As a group, PFL1 neurons tile the PB and FB. We observed the tiling pattern of PFL1 is mostly identical across the two electron microscopy datasets we analyzed, although we found a small difference in the PB (Extended Data Figure 1a-c); this difference, however, does not meaningfully impact the result of our model simulations described below (Extended Data Figure 1d-f). In the PB, PFL1 neurons are postsynaptic to the EPG (’compass’) neurons, which compute the fly’s heading, and Δ7 neurons, which smooth the fly’s heading to two sinusoidal profiles^23,32^. In the FB, the dendrites of PFL1 neurons interleave (Supplementary Video 1) and receive input from FB neurons. Finally, PFL1 neurons send axons unilaterally to either the left or right lateral accessory lobe (LAL), a premotor area^51^.

**Figure 1:**
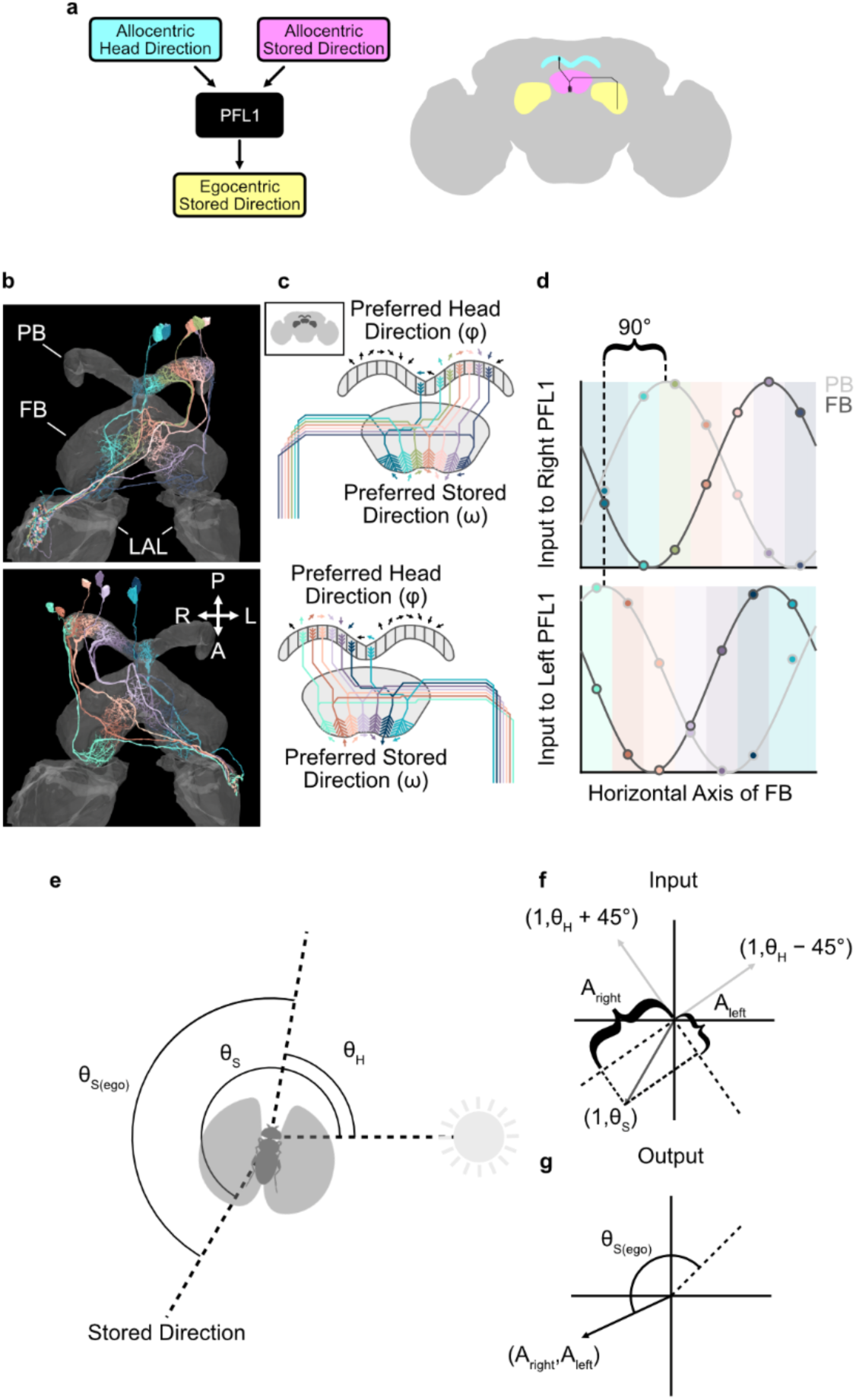
Anatomically constrained model of PFL1. **a,** We hypothesize that PFL1 takes as inputs the allocentric head direction and an allocentric stored direction and then outputs the egocentric stored direction. **b,** Rendering of PFL1 neurons from hemibrain dataset. Top: neurons with axons on the right side of the brain. Bottom: Neurons with axons on the left side. P, posterior. L, left. A, anterior. R, right. **c,** Drawing of PFL1 neurons that illustrates how they tile the PB and FB and how these tiling patterns define each neuron’s preferred head direction (φ) and preferred stored direction (ω). Inset: drawing of brain with PB, FB, and LAL. **d,** Graph showing the simulated input to PFL1 neurons for the value of head and stored directions shown in e. Each dot represents the input to a neuron. The color of the outline of the dot indicates whether the input is from the PB or FB. Light grey indicates the activity in the PB. Dark grey indicates the hypothetical activity in the FB. The x-axis is the horizontal axis of the fan-shaped body. The shaded regions indicate the portion of the fan-shaped body innervated by each neuron. The brace denotes that the PB input to neurons with axons on opposite sides of the brain is 90° out of phase. **e,** Drawing showing the meaning of the variables from the model: head direction (θ_H_), stored direction (θ_S_), and egocentric stored direction (θ_S(ego)_). **f,** Graph showing the input to PFL1 as vectors for the values of θ_H_ and θ_S_ shown in e. The braces denote the output of the model (A_right_ and A_Left_). The coordinates in this graph are polar coordinates. **g,** Graph showing the output of the model (A_right_ and A_Left_) as a vector for the values of θ_H_ and θ_S_ shown in e. The set of coordinates in this graph are Cartesian coordinates. The dotted line is the direction of the vector when θ_H_ and θ_S_ are equal. The difference between this line and the direction of the vector is equal to the egocentric stored direction (θ_S(ego)_).

### The anatomy of PFL1 neurons provides a computational scaffold for coordinate transformation

To help predict PFL1’s function, we built a circuit model constrained by its anatomy. We used the well-characterized head direction tuning of the EPG and Δ7 neurons, which is based on both topographical arrangement of these neurons and their physiology, to predict the input to each PFL1 neuron in the PB and define its preferred head direction (Fig. 1b). Since Δ7 neurons exhibit a sinusoidal activity profile across the PB^23,32^, we modelled the input that each PFL1 neuron receives in the PB as a sinusoidal function of the difference between the fly’s current head direction and the neuron’s preferred head direction. In the FB, on the other hand, the function and the activity profile of the FB neurons presynaptic to PFL1 have not been identified. Extrapolating from known properties of PFL2 and PFL3, which have similar morphologies with PFL1 and receive goal direction information from presynaptic neurons in the FB^25,46^, we predicted that FB neurons presynaptic to PFL1 encode a vector with an arbitrary direction defined in an allocentric reference frame. We call this the *stored direction,* and we speculate it is encoded by the phase of sinusoidal input to PFL1’s FB dendrites. Thus, we modeled the input to PFL1’s FB dendrites as a sinusoidal function of the difference between each neuron’s preferred stored direction––determined by its position in the FB––and the stored direction (Fig. 1c).

A key feature of PFL1 anatomy is the pattern of dendritic projections to the PB and the FB. PFL1 neurons with axons in the left LAL project their dendrites to the FB in the opposite direction from neurons with axons in the right LAL (Fig. 1b). This pattern causes left and right PFL1 neurons with dendrites located in a similar part of the FB to have preferred head directions separated by 90° (Fig. 1c and Extended Data Fig 2a). Thus, this pattern can be thought of as inducing a ±45° shift in the phase of the head direction input. As a result of this anatomical feature, two identical sinusoidal population activity profiles of the head direction signal from each side of the PB become 90° out of phase when they meet information from the FB (Fig. 1d and Extended Data Fig. 2b). Conceptually, two identical copies of a head direction vector are rotated ±45° in PFL1 neurons, forming a 90° angle between them when they receive the stored direction vector input from the FB.

In the LAL, downstream neurons pool input from all PFL1 neurons with axons on the same side of the brain (Extended Data Fig. 3). Thus, we modeled the output of PFL1 as two numbers reflecting the sum of the output of all the PFL1 neurons on the right or left side of the brain.

Taking everything together, PFL1 is divided into two populations (left and right), which both receive two vectors as input and each generate a single scalar value as output. This network structure is poised to calculate the inner product between each population’s input vectors. To implement this function, the output of each PFL1 neuron would have to be equal to the product of its two inputs (signed multiplication; Extended Data Fig. 4), which is an important and the only computational assumption we make in our model. Through simulations (Extended Data Fig. 5) and analysis (mathematical appendix), we also found that models using two other nonlinear activation functions, ‘unsigned multiplication’ and ‘squares of the sum’, produce nearly identical outputs to that of a model using signed multiplication (Mathematical Appendix). Thus, we opted to use the signed multiplication model in the rest of this study.

If PFL1 calculates the inner product between its input vectors, mathematically this is identical to calculating projections (braces in Fig. 1f) of the stored direction vector (dark grey vector in Fig. 1f) onto two orthogonal vectors that are ±45° rotated versions of the head direction vector (light grey vectors in Fig. 1f). This results in two coordinates (A_left_ and A_right_ in Fig. 1g) that define an output vector that encodes the stored direction in an egocentric reference frame (8_s(ego)_ in Fig. 1g). Thus, our modeling suggests a possible function of the PFL1 neurons: They transform an allocentric vector stored in the FB to an egocentric reference frame by comparing it to the head direction information (Fig. 1a).

### Input-output relationship of PFL1 is consistent with the coordinate transformation model

The coordinate transformation model described above makes three predictions about PFL1’s input-output relationship: (i) the output of PFL1 should have a sinusoidal relationship with the head direction, (ii) the phases of the outputs in the left and the right LALs should be 90° apart, and (iii) the phase of the output relative to the head direction should be determined by the stored direction (Extended Data Figure 4a). We tested whether PFL1’s physiology is consistent with these predictions. To simultaneously record the head direction input to PFL1 as well as PFL1’s output, we expressed the glutamate sensor iGluSnFR3^52^ and the red-shifted calcium indicator jRGECO1a^53^ in PFL1 and performed 2-photon imaging in head-fixed flies flying in a virtual reality environment (Fig. 2a). The iGluSnFR3 signal in the PB indicates glutamatergic input from Δ7 neurons representing the head direction. Note, since the glutamatergic input to PFL1 from Δ7 is inhibitory, the head direction (defined as the phase of the positive PB input) is equal to the population vector of the iGluSnFR3 signal in the PB plus 180°. The jRGECO1a signal in PFL1 axons in the LAL reports the sum of PFL1 neurons’ output on each side.

**Figure 2:**
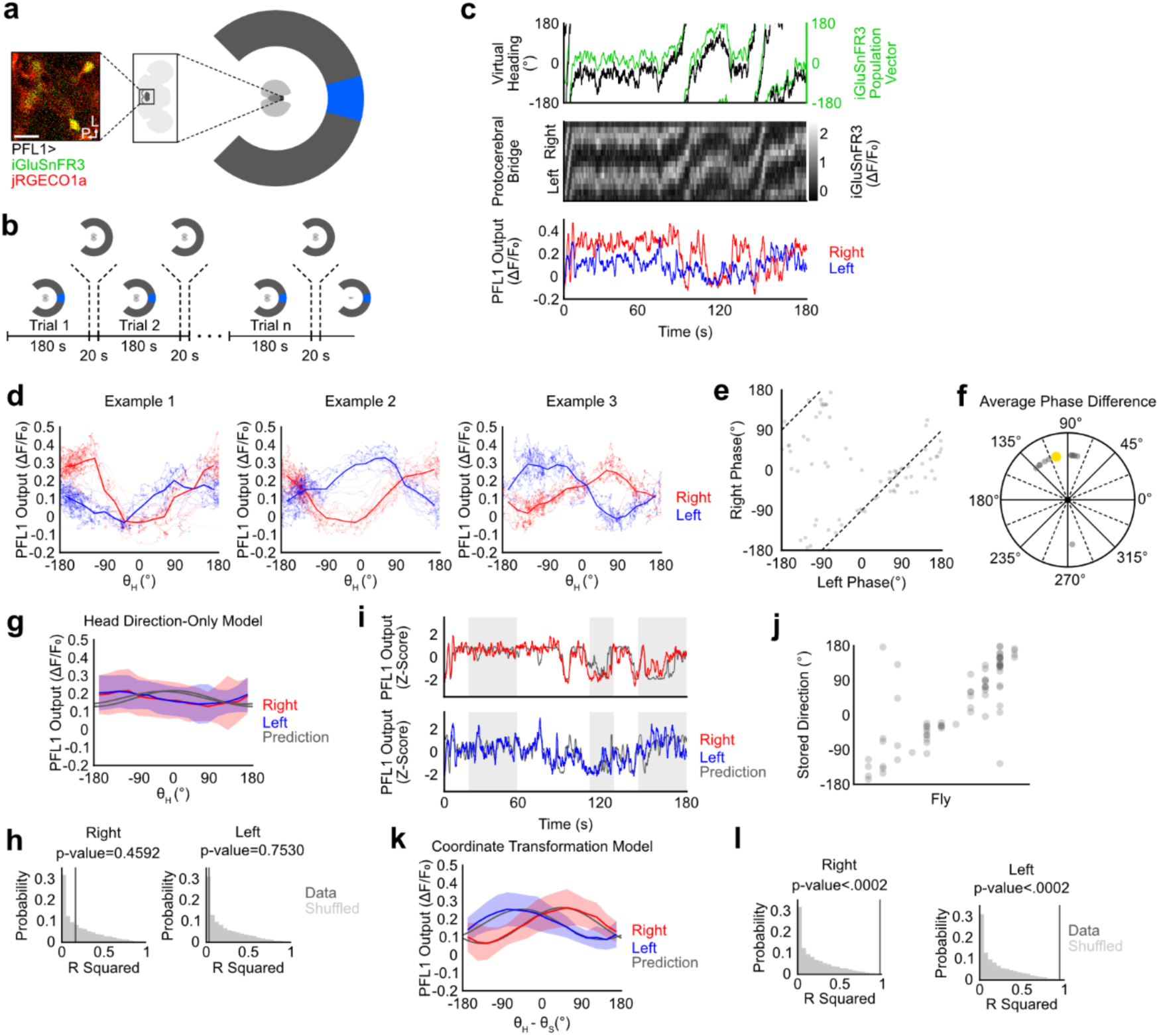
Inferring stored direction from PFL1 input output relationship. **a,** A schematic illustrating the experiment. Scale Bar 20 micrometers (µm). L, left. P, posterior. **b,** Trial structure of experiment. **c,** Data time series from an example trial. **d,** Examples showing the PFL1 output as the function of head direction. Each dot is a time point. Thick lines are the averages of the dots. Example 1 is the same data from c. **e,** A dot is plotted for each trial based on the phase for the left and right, found by fitting a cosine function to the entire trial. Dotted line indicates a 90° phase difference between the left and right. **f,** The average phase difference between the cosines fit to the left and right for each trial (left phase - right phase). Grey dots are single-fly averages. The yellow dot is the across-fly average, 104.9°. **g,** Output of PFL1 plotted as a function of the head direction. Blue and red lines are across-fly averages. The shaded region represents the standard deviation. N =11 flies. Grey lines are predictions based on the head direction-only model. **h,** R-squared values between the head direction-only model and across-fly averages for data (dark grey line) and shuffled data (light grey distribution). **i,** Example showing the model fit to the data with our cross-validation method. The grey traces are simulations from the model. Blue and red traces are data. The time intervals with a white background were used for fitting. Time periods with grey backgrounds were held out for testing. These time periods were selected randomly for each trial. **j,** The stored direction for each trial, found through fitting. Each dot is a trial, and each column is a fly. The x-axis is sorted based on the average stored direction. **k,** The output of PFL1 plotted as a function of the difference between the head direction and stored direction. This x-axis aligns the data according to the fitted stored direction. Red and blue lines are across the fly averages. The shaded region represents the standard deviation. N =11 flies. Grey lines are the prediction from the coordinate transformation model. **l.** The same as f for the coordinate transformation model.

We divided each experiment into 3-minute trials punctuated by 20 seconds of darkness (Fig. 2b). As expected, the iGluSnFR3 signal in the PB reliably tracks the fly’s heading in the arena (Fig. 2c). Plotting the axonal activity of PFL1 as a function of the head direction reveals a sinusoidal relationship (Fig. 2d and Extended Data Fig. 6a-b). Critically, the right and left outputs in LAL are approximately 90° out of phase from each other (Fig. 2e,f and Extended Data Fig. 6c,d,), as our anatomically-constrained computational model predicts (Extended Data Fig. 4a).

The phase of the PFL1 output relative to the head direction varies between flies (Fig. 2d,e). This cannot be explained if the PFL1 output is only influenced by the head direction information. Furthermore, pooling data across flies does not match a prediction based on a head direction-only model (Fig. 2g,h), strongly suggesting that an additional input, likely from the FB, is influencing the PFL1 output. As we introduced above, we call this information from the FB a stored direction.

We fit the model to infer the stored direction encoded in the FB using the head direction in the PB and the PFL1 output in the LAL for each trial. We did this fitting and simulation using a cross-validation technique (Fig. 2i). We found that the best-fit stored direction tends to be consistent across trials for the same fly, but appears arbitrary across flies (Fig. 2j), important evidence that the stored direction depends on an individual’s distinct experience. Our physiology experiments also showed that it can sometimes abruptly change within the same fly (Extended Data Fig. 7), suggesting that it is flexible.

Finally, when we aligned the data based on the best-fit stored direction prior to averaging across flies, we found a close match between the data and a prediction based on the coordinate transformation model (Fig. 2k,l). Together, these results show that the output of PFL1 resembles a set of Cartesian coordinates that encode the stored direction in an egocentric reference frame. Overall, PFL1’s output has three physiological features: (i) it has a sinusoidal relationship with the head direction, (ii) the phases of the outputs in the left and the right LALs are 90° apart, and (iii) the phase of the output relative to the head direction is flexible.

### The PFL1 model provides a powerful framework to investigate mechanisms encoding the stored direction

The coordinate transformation model predicts that the phase of the sinusoidal input in the FB sets the phase between the output of PFL1 and the head direction. Thus, we sought to use our ability to infer the stored direction from the PFL1 input-output relationship to dissect the function of the circuitry upstream of PFL1 in the fan-shaped body.

To this end, we used the electron microscopy connectome data to canvass FB neuron types presynaptic to PFL1. We found that FC1 neurons provide the largest input to PFL1 in the FB by synapse count (Fig. 3a and Extended Data Fig. 8)^18^. Because of this strong input to PFL1, we hypothesized that FC1 may encode the stored direction and provide this information to PFL1. On the other hand, FC1 receives input from the PFNa neurons, which convey head direction information to the FB^28,54^. Thus, another reasonable hypothesis is that FC1 might display head direction tuning. To test these two mutually exclusive hypotheses, we simultaneously recorded the activity of FC1, Δ7, and PFL1 by expressing jRGECO1a in FC1 and Δ7 and GCaMP7f in PFL1^55^. Note that the split gal4 line labels only a subset of FC1, limiting our ability to test the hypotheses for the entire FC1 population. We fit the coordinate transformation model to the Δ7 and PFL1 data to find the best-fit stored direction for each trial (Fig. 3b). If FC1 encodes the stored direction, the phase of calcium activity across the width of the FB should be equal to the stored direction we find through fitting the model to the Δ7 and PFL1 recordings. We found, however, that the phase of the FC1 activity did not match the stored direction (Fig. 3c,d,f,g). Instead, it tracks the head direction (Fig. 3c,e-g). Therefore, FC1 calcium activity seems to reflect its input from PFNa rather than the stored direction that we infer PFL1 receives in the FB.

**Figure 3:**
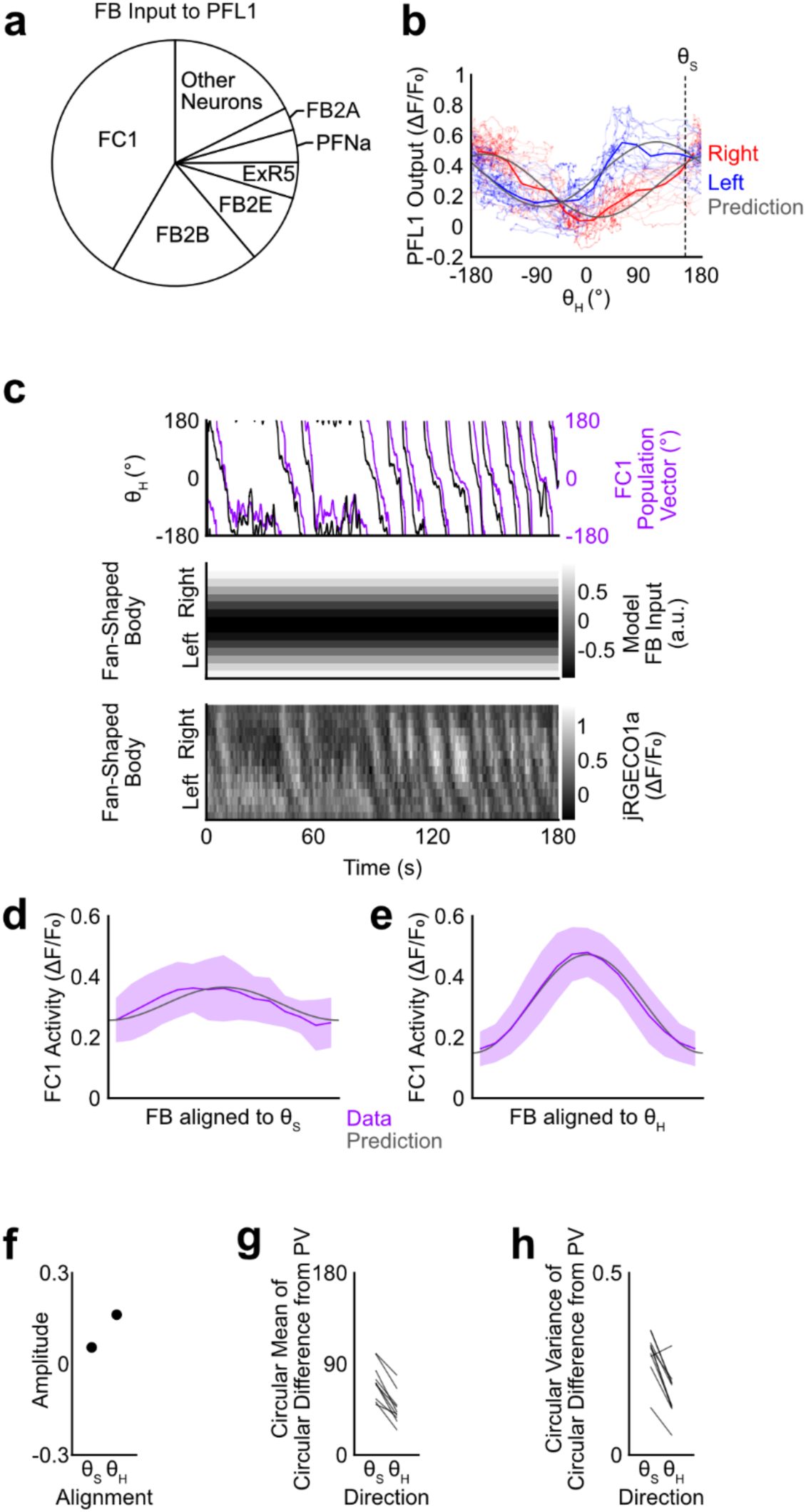
Simultaneous recording of Δ7, FC1, and PFL1. **a,** A pie chart showing proportion of synapses from each cell type presynaptic to PFL1 in the Fan-shaped Body, by synapse count. **b,** Data from example trial with prediction from the coordinate transformation model (grey lines) based on a stored direction found through fitting (dotted line). Each dot is a time point. Red and blue lines are averages of the same color dots. **c,** Data time series from same trial as b. Top: Head direction (black) and FC1 population vector (purple). Middle: Predicted FB input from coordinate transformation model. Bottom: Measured FC1 activity. **d,** FC1 activity plotted as a function of the position in the Fan-shaped Body aligned to the stored direction. Purple line is an across-the-fly mean. The shaded region represents the standard deviation. The dark grey line shows the predicted result if the phase were to encode the stored direction with an amplitude fit to the data. N = 10 flies. **e,** The same as d except aligned to the head direction. **f,** The amplitude found by fitting the prediction. **g,** The circular mean of the circular difference between the FC1 population vector and the head direction or the stored direction. **h,** The circular variance of the circular difference between the FC1 population vector and the head direction or the stored direction. P-value = 0.0039, Wilcoxon sign rank test

The second largest input to PFL1 in the FB is a cell type called FB2B (Fig. 3a). FB2B is a type of tangential neuron that spans the width of the FB (Extended Data Fig. 8c). If the calcium activity reflects the overall activity of the neuron, we would expect calcium to be uniform across the width of the FB. However, a sinusoidal pattern of calcium activity in the axons of FB2B might be a mechanism for providing the stored direction input to PFL1. When we simultaneously recorded calcium in FB2B, Δ7, and PFL1, the calcium signal of FB2B was uniform across the width of the FB (Extended Data Fig. 9a,b). Aligning the FB2B data to either the stored direction or the head direction results in an amplitude close to zero, suggesting it does not have a meaningful phase in its calcium activity (Extended Data Fig. 9c-f). Overall, these results constrain possible mechanisms for encoding a stored direction in the FB and providing this information to PFL1.

### PFL1 is involved in steering maneuvers

PFL neurons (and their homologs in other insects) are generally thought to control steering, because their axons project to the LAL, a premotor area^19,21^. We also noticed in our calcium imaging experiments that PFL1 activity declined whenever a fly stopped flying (Extended Data Fig. 10). This made us wonder if PFL1 might control steering during flight.

In our imaging experiments, where we infer a constant stored direction from PFL1’s input output relationship, flies are not engaged in goal-directed locomotion. For example, in a fly that is flying in circles, PFL1 still reads out a constant stored direction (Fig. 4a,b). In fact, most flies we recorded from did not show a preference to fly towards the stored direction (Fig. 4c). This suggests that the stored direction that PFL1 transforms and conveys to downstream neurons only specifies a goal in a specific behavioral context not present in this experiment. This would be in line with EPG silencing experiments where the CX is not essential for all navigational behaviors, such as object fixation^43,56^.

**Figure 4:**
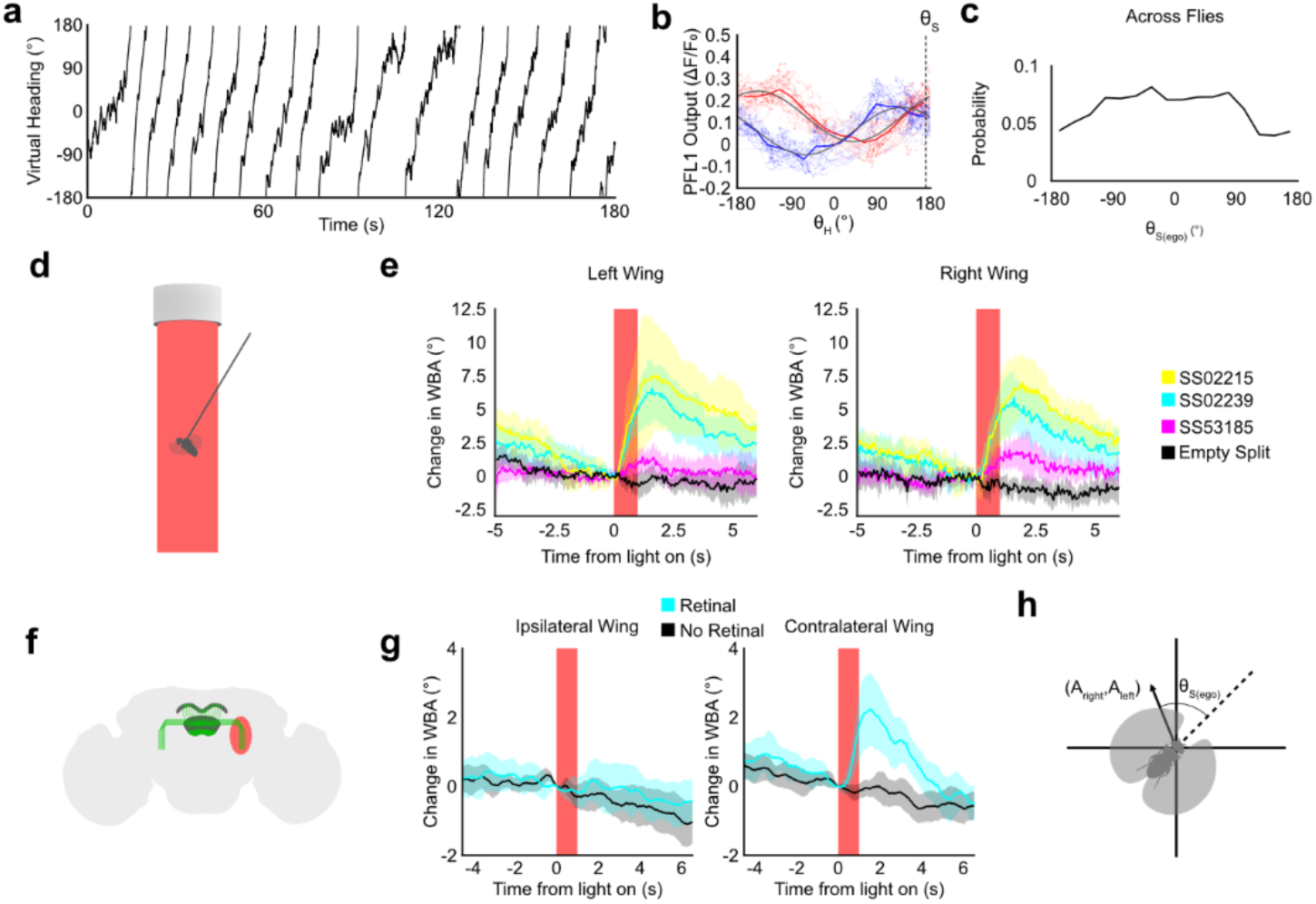
Probing the behavioral role of PFL1 activity. **a,** The behavioral trajectory of an example fly that was flying in circles. **b,** PFL1 output plotted as a function of head direction for the example fly from a. Grey lines are the prediction from the coordinate transformation model based on a stored direction found through fitting (dotted line). Each dot is a time point. Red and blue lines are averages of the same color dots. **c,** Histogram showing the egocentric stored direction across flies. If flies preferred to fly in the stored direction, there would be a peak at 0. N=11 flies. **d,** A schematic showing illumination of an entire fly with red light. **e,** A change in wing beat amplitude upon bilateral PFL1 stimulation. The shaded region represents the 95% confidence interval. SS02215 N=6 flies. SS02239 N=6 flies. SS53185 N=6 flies. Empty Split N=7 flies. **f,** A schematic showing unilateral red-light stimulation using a digital micromirror device. **g,** A change in wing beat amplitude upon unilateral PFL1 stimulation. The shaded region is the 95% confidence interval. N = 10 flies for retinal. N=10 flies for no retinal. **h,** A schematic showing the relationship between PFL1 output (A_right_, A_left_) and the egocentric stored direction (θ_S(ego)_).

To test if PFL1 influences locomotion, we optogenetically activated the neurons. Although a previous study found that activating PFL1 neurons in walking flies does not induce any steering behavior^25^, we reasoned based on PFL1 neurons’ activity in flying flies that flight may be the appropriate behavioral context. First, we bilaterally activated PFL1 neurons in tethered flying flies by expressing CsChrimson in PFL1 using three different split gal4 lines and pulsed red light on the entire fly. (The flies were rendered blind by a mutation in the *norpA* gene to eliminate any visual response to light.) Bilateral activation of PFL1 induced a bilateral increase in wing beat amplitude (Fig. 4d,e).

Having established that PFL1 activation induces a behavioral response during flight, we posited that activating PFL1 neurons in only one hemisphere should generate a steering response. To unilaterally activate PFL1 neurons, we used a digital micromirror device to expose the axons of PFL1 on only one side of the brain to red light. We observed an increase in the contralateral wing beat amplitude (Fig. 4f,g). Such a change in wing beat amplitude would drive an ipsilateral turn in a freely flying fly.

In the context of our model, stimulating the PFL1 neurons in the right LAL would increase one component of the output vector (the X-coordinate in Fig. 4h), signaling that the egocentric stored direction is to the fly’s right, while stimulating the left neurons would signal the egocentric stored direction is to the left (Fig. 4h). The behavioral response to these stimulations is to turn towards the egocentric stored direction.

Overall, these results suggest that the stored direction may encode a goal direction and that PFL1 neurons provide an egocentric steering signal that the downstream motor circuits can use to turn the animal towards the goal. Yet, animals do not always use it to adjust their locomotion, suggesting that it depends on behavioral contexts.

## Discussion

When navigating, the brain generates and stores many types of information––head direction, grid, place, border, and goal-related signals––that it uses to make navigational decisions. Many of these signals are distinct from motor commands: Indeed, many do not have immediate motor implications and instead encode aspects of the animal’s navigational *state* rather than, say, a turning decision.

Here, we describe an example of how the brain combines multiple navigational signals to create new signals: PFL1 neurons innervating multiple brain areas (PB, FB, and LAL) integrate the sensory-driven head direction signal and a stored, allocentric vector by performing a coordinate transformation to generate a new, egocentric vector that can be used to control navigation. The most striking result is how well the predictions from the anatomically constrained models match the physiological observations. The dendritic innervation of PFL1 neurons in the PB, the FB, and the outputs on each side of the LAL are patterned in a highly organized way, which allowed us to develop computational models with minimal assumptions.

This enabled us to develop a computational model that shows how PFL1 can perform a vector computation that would be useful for navigation. This model provided a framework for interpreting our physiological recordings of PFL1. The fact that the physiological recordings of PFL1 neurons match the predictions of the model testifies to the power of using EM-based connectivity information in discovering the dynamics of neural networks and their functions.

Our characterization of the vector stored in the FB that provides part of PFL1’s input suggests it may encode a flexible goal: Our physiological experiments in PFL1 showed that it is maintained for an extended period of time (across trials that span tens of minutes), but it is also flexible (across individuals and some timescales in some flies), presumably representing fly’s distinct experience. This simultaneous exhibition of both stability (across some timescales) and flexibility (across individuals and some timescales) is also a characteristic of the mapping of the orientation of a visual scene to the head direction representation encoded by the EPG neurons^57,58^, suggesting that such flexible yet stable navigational signals may be common in the fly central complex.

Our model suggests that the phase of the PFL1’s input-output relationship should reflect the direction of a vector encoded by the input PFL1 receives in the FB. Although we found that the calcium activity of FC1 neurons and FB2B neurons––PFL1 neurons’ primary synaptic partners in the FB––does not encode this vector, it remains possible that they supply the vector through an alternate mechanism. For example, plasticity of tangential neuron synapses may encode vectors^43,44^. For this to work, the synaptic weights of tangential neurons would be reshaped when a salient event occurs in a pattern that reflects the activity of upstream columnar neurons. This is remarkably similar to structural and computational mechanisms regulating the synapses between ER neurons (tangential in the EB) and columnar EPG neurons that encode the head direction^57,58^. Applying the same logic to the circuitry upstream of PFL1, the synaptic connections from FB2B or other tangential neurons to columnar PFL1 neurons may store a vector, which would help store the directional information on longer timescale than calcium activity. Testing this hypothesis, however, requires a reliable method to measure the synaptic weights between FB2B and individual PFL1 neurons, which remains a technical challenge at this point.

PFL1 neurons perform a unique function among PFL neurons. PFL1 differs anatomically from PFL2 and PFL3: its FB presynaptic partners are different than PFL2 and PFL3, and PFL1 innervates a separate LAL region from PFL2 and PFL3^25,46^, suggesting distinct vectorial inputs and motor-related outputs^54^. Furthermore, unlike PFL3, which shows 135° phase differences in LAL and thus functions well for steering left or right^25,46^, PFL1’s 90° difference is more suited to encoding a vector in a two-dimensional Cartesian coordinate system. Operationally, it may provide an egocentric direction to a certain location, such as a home vector or a vector to a recently encountered food location, which needs to be tracked while navigating out of such locations.

Going forward, the PFL1 circuit provides a powerful entry point to uncover how internally maintained and sensory-driven information is integrated to shape flexible navigational behavior. Our ability to simultaneously record the head direction input to PFL1 and the output of PFL1 to infer the stored direction in behaving flies while also recording or perturbing a third cell type in the fan-shaped body will allow us to decipher how the stored direction is formed and identify behavioral contexts that gate the impact of this vector on motor decisions. Because head direction cells are ubiquitous across the animal kingdom, understanding how the PFL1 circuit uses the fly’s internal compass to relate information stored in an allocentric reference frame to its immediate surroundings will yield insights that generalize well beyond the fly’s small brain.

## Supporting information

Anatomy of PFL1

## Acknowledgments

This study was supported by the National Eye Institute of the National Institutes of Health (DP2EY032737; 1U01NS131438-01). The content is solely the responsibility of the authors and does not necessarily represent the official views of the National Institutes of Health. It was also supported by Air Force Office of Scientific Research (FA9550-23-1-0722) and National Science Foundation (2440847), Searle Scholars Program, Sloan Research Fellowship, Klingenstein-Simons Fellowship in Neuroscience, and McKnight Foundation. We thank T. Wolff and G. Rubin for sharing unpublished split Gal4 lines, D. Bushey for sharing the iGluSnFR3 flies, the Simpson lab for sharing their space while our lab was setting up, SciDraw.io, G. Costa, and A. Bates for vector graphics files of a fly (DOI: 10.5281/zenodo.3926137) and a fly brain (DOI: 10.5281/zenodo.4421155), T. Crahan, E. Chen, and D. D’Lima for experimental assistance, O. Vera and E. Jung for technical support, and J.-M. Knapp for valuable discussions and comments on the manuscript.

## Author Contributions

BG and SK conceived the study. BG collected and analyzed data. BG and SK wrote the manuscript.

## Competing interests

The authors declare no competing interests.

## Data availability

All raw physiological data will be available upon request. All processed data is available at public repository by the time of publication.

## Code availability

Analysis code will be available by the time of publication.

## Methods

### Model

#### Coordinate Transformation Circuit Model

We modeled the input to each PFL1 neuron in the PB/FB as a cosine function of the head/stored direction minus the neuron’s preferred head/stored direction, which is determined by their horizontal location in each area. In our model, the output of each PFL1 neuron is the product of its two inputs, as in the following equation.

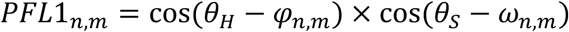

where *PFL1_n,m_* is the output of an individual PFL1 neuron, with *n* representing an index of a PFL1 neuron and *m* representing where the axon is (either the left or the right side of the brain). *θ_H_* is the animal’s head direction and θ_S_ is the stored direction in the FB. *φ _n,m_* and *ω _n,m_* represent the preferred head direction and the preferred stored direction of *PFL1_n,m_*, respectively. We defined *φ* to be [−112.5 −67.5 −22.5 22.5 67.5 112.5 180] for *m = right* and [157.5 −135 −90 −45 0 45 90] for *m = left*. We defined *ω* to be [−168 −120 −72 −24 24 72 132] for *m = right* and [−156 −96 −48 0 48 96 144] for *m = left*. For the flywire model in Extended Data Fig. 1, we used the following values of *φ*: [−112.5 −67.5 −22.5 22.5 67.5 112.5 168.75] for *m = right* and [168.75 −135 −90 −45 0 45 90] for *m = left*. Both models resulted in nearly identical results.

LAL neurons downstream of PFL1 sum the input of all the PFL1 neurons with axons on the same side of the brain. Thus, the final output of the model, A, is the sum of PFL1 neurons in each side.

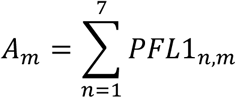

#### Head Direction-Only Model

For the head direction only model, we set the output of each PFL1 neuron to be equal to its protocerebral bridge input, but otherwise the model was the same.

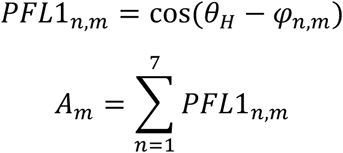

#### Alternative Activation Functions

For models with different activation functions, we changed the equation describing the output of each PFL1 neuron. For unsigned multiplication, we added 1 to the cosine functions.

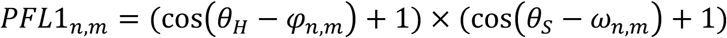

For the square of sum model, we used the following equation.

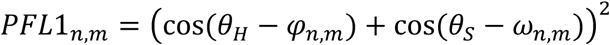

#### Fly Rearing

We raised flies on a standard Würzburg recipe, collected them daily, and used them 7 days after collection. We used female flies for all two-photon experiments. For our optogenetic experiments, we used flies with a mutation in the *norpA* gene, which is on the *X* chromosome. Since it is easier to generate males with this mutation because they are hemizygous for *X*, we used male flies for our bilateral optogenetic experiments. Male flies do not fly as well as female flies, however. Therefore, for our unilateral optogenetic experiments, we used female flies. See Table 1 for the genotype of the flies used in each experiment.

#### Two Photon Imaging

We cold anesthetized flies, removed their front legs to prevent them from pushing the metal holder, waxed their proboscis in place, and tethered flies to a pin using UV curable glue; then, we removed them from cold. We glued the head to the bottom of an inverted stainless-steel pyramid with a small opening at the tip. After filling the pyramid with saline, we removed the cuticle and trachea to expose the brain. Then we placed them in the center of a 270° LED arena (https://reiserlab.github.io/Modular-LED-Display/G3/) under the objective of a custom two-photon microscope (https://www.janelia.org/open-science/mimms-22-2024)^59,60^. We covered the surface of the LED arena with gel filters to avoid light leakage to PMTs (Roscolux Cerulean Blue R375 and Congo Blue R382), and we added a layer of paper on top of the filters to create a matte surface to prevent reflections in the arena. We tracked the wings in real time using a camera and custom wing tracking software as described previously^33^, to calculate the difference between left and right wing stroke amplitudes. We used it to control the angular speed of the scene, which contained a single blue bar, in all of our closed-loop experiments.

For two-color imaging, we used a long wavelength of a tunable two-photon laser. We explored and chose the best wavelength for each experiment: 1030nm for the iGluSnFR3 and jRGECO1a experiments in Fig. 2 and 4 and Extended Data Fig. 6 and 7; 1000 nm for the Δ7, PFL1, and FC1 imaging experiments in Fig. 3; 1020nm for the Δ7, PFL1, and FB2B imaging experiments in Extended Data Figure 9. We used 930nm for imaging of PFL1 in Extended Data Figure 10.

### Optogenetic Stimulation

#### Bilateral Stimulation Experiments

We cold anesthetized flies, tethered them to a pin using UV curable glue, and mounted them under a collimator attached to a 625 nm LED via an optic fiber. We tracked wing beat amplitude with a camera, an IR LED for illumination, and custom software. We presented red light in a pulse train of five 1-second pulses interspersed with 10-second interpulse intervals. A fly had to fly for an entire pulse train for the data to be included in the analysis. If flies stopped flying, we applied a short air puff to induce flight.

#### Unilateral Stimulation

We prepared flies as those for two-photon imaging and mounted them under the two-photon microscope, which we briefly used to locate the PFL1 neurons. We used custom software to draw regions of interest (or masks) around the axons of PFL1 to be optogenetically stimulated. We aligned a digital micromirror device (DMD) to the light path of the microscope and delivered a 617 nm light (Thor Labs Part Number: M617L3) to the DMD via a liquid light guide. We stacked two longpass filters (Semrock Part Number: BLP01-594R) and placed them in front of the output of the DMD to block wavelengths leaking to PMTs. For these experiments, we used a 561 nm single edge dichroic mirror (Semrock Part Number: Di03-R561) to allow the red light from the DMD and the two-photon laser to pass through while reflecting emitted light from the sample to the green PMT. We induced flies to fly and presented 1-second LED pulses of light every 10 seconds while tracking the flies’ wings. The two-photon laser was turned off during optogenetic stimulation. We switched the masks between activating the PFL1 axons on the left side and the right side of the brain. We randomized the order of these stimulations between flies. We looped the stimulations until the fly would no longer fly.

### Analysis

#### Image Processing

We predetermined the background noise level by measuring the oscillatory noise from the PMT. We then subtracted this level from all imaging data and half-rectified the data before further analysis. We divided the PB into either 14 ROIs for experiments where PFL1 expressed iGluSnFR3 or 18 ROIs for experiments where Δ7 expressed jRGECO1a to approximate the PB glomeruli. We divided the FB into 15 ROIs horizontally to most closely match the tiling pattern of the PFL1 dendrites. We drew separate polygons around the left and right LAL to define their ROIs. A running average intensity projection of a volume (five planes) at a given time was computed for each pixel. We applied an approximately 1-second rolling average to the data to smooth the data. To calculate the ΔF/F_0_, we defined the F_0_ for each ROI as the tenth percentile frame for each ROI after smoothing.

#### Calculating Population Vector

To calculate population vectors for the PB and the FB, we assigned each ROI a vector with a direction based on its position in the neuropil and a length equal to the ΔF/F_0_ at that time point. Then, we calculate the average of these vectors to calculate the population vector.

#### Fitting Model

For fitting the circuit model to the data in Fig. 2, we divided each trial into 10 time bins. We randomly selected half of the time bins to use for fitting the model. We left the other half out for testing the model. For every time point, we defined θ_H_ as the PB population vector, calculated from iGluSnFR3 signal, plus 180 degrees to account for the inhibitory nature of Δ7 neurons. We used Matlab’s fminbnd function to find the value of θ_S_ that maximize Pearson’s correlation coefficient between the model prediction and the jRGECO1a signal in PFL1’s axons on both sides.

We used the same fitting process in Extended Data Fig. 7. However, we did not break the experiment up into trials, so we manually determined the time periods over which to fit. From visual inspection, it appeared that the difference between the direction of a vector defined by the PFL1 calcium recordings and the head direction changed at a certain point in the experiment. We selected time intervals immediately before and after that time point to fit the model. Unlike Figure 2, we left no data from those time periods out of the fitting process, because the model was not quantitatively compared to the data.

For Fig. 3 and Extended Data Fig. 9, we used a similar fitting process to find an optimal value of θ_S_ for each trial, given the Δ7 recordings in the PB and the PFL1 recordings in the LAL. However, we used data from the entirety of each trial to fit the model, because the result of the fitting was being compared with the FB data, which was not used to fit the model.

#### Shuffling

For the histograms in Fig. 2 and Extended Data Fig. 6, we generated null distributions for a given test statistic based on chance. We circularly permutated the LAL recordings from each trial by a random interval before performing the analysis. The shuffling and analysis were repeated 5000 times to generate the null distributions.

#### Cosine Fitting

In Fig. 2e-f, and Extended Data Fig. 6, we fit a cosine function to the recordings of PFL1 activity in the LALs. We used Matlab’s fminsearch function to find the optimal amplitude, phase, and offset for a cosine function of the head direction that minimized the root mean square error between the cosine function and the PFL1 recording for every time point in a trial. To avoid fitting and testing with the same data, we divided trials into 10 time bins and randomly selected half to use for fitting and half for testing (Extended Data Fig. 6 A-B). For analysis where data was not directly compared to a cosine function fit to the same data, we used the entire trial to fit the cosine function (Fig. 2e-f, Extended Data Fig. 6 C-D).

#### Aligning FB to Head and Stored directions

To generate the plots in Fig. 3d, e and Extended Data Fig. 9c, d, we parameterized the position in the FB by assigning each FB ROI an angle. We assigned the data for each ROI for each time point an x coordinate based on the difference between θ_S_ or θ_H_ and the angle assigned to each ROI. We divided the x-axis into 16 bins and calculated the mean of all the time points within each bin for each trial. Then we averaged the bins across all the trials for each fly. We calculated the mean and standard deviation for each bin across flies. We fit a prediction to the across fly mean using Matlab’s fitlm function. The prediction is equal to the cosine of the x-axis.

#### Electron microscopy analysis

To find the cell types presynaptic to PFL1 in the FB for Fig. 3a and plot the connectivity matrix in Extended Data Fig. 8, we used neuPRINT’s (https://neuprint.janelia.org/) custom query feature to download a list of PFL1’s postsynapsic sites in the FB.

To create the connectivity matrix in Extended Data Fig. 1c (bottom), we used neuPRINT to download all the connections between EPG neurons and PFL1 neurons.

To create the connectivity matrices in Extended Data Fig. 1c (top) and Extended Data Fig 3, we used the Connectivity Matrix widget in CATMAID (https://fafb-flywire.catmaid.org/).

To create the images in Fig. 1a, Extended Data Fig. 1a-b, and Extended Data Fig. 8b-c, meshes of the neurons were downloaded using CloudVolume (https://github.com/seung-lab/cloud-volume), and the images were rendered in Blender.

## Genotype Table

Genotype Table:
| Figure | Genotype |
| --- | --- |
| 2, 4a-c, Extended Data 6, Extended Data 7 | (female) UAS-iGluSnFr3v857-PDGFR (attP18);;UAS-jRGECO1a x (male) SS02239 |
| 3 | (female) 84H05-LexA (attP40), UAS-jRGECO1a (su(Hw)attP5);SS60353 x (male) 55G08-Gal4 (attP2) LexAop-GCaMP7f (VK00005) |
| 4e | (female) w<sup>+</sup>,norpA<sup>36</sup>,UAS-CsChrimson-mVenus (attP18) x (male) SS02215 or SS02239 or SS53185 or empty split |
| 4g | (female) w<sup>+</sup>,norpA<sup>36</sup>,UAS-CsChrimson-mVenus (attP18) x (male) w<sup>+</sup>,norpA<sup>36</sup>;SS02239 |
| Extended Data 9 | (female) 84H05-LexA (attP40), UAS-jRGECO1a (su(Hw)attP5);49H05-Gal4 (attP2) x (male) 55G08-Gal4 (attP2) LexAop-GCaMP7f (VK00005) |
| Extended Data Fig. 10 | (female) UAS-GCaMP6f (VK00005), UAS-tdTomato (attP2) x (male) SS02239 |

---

**Extended Data Figure 1:**
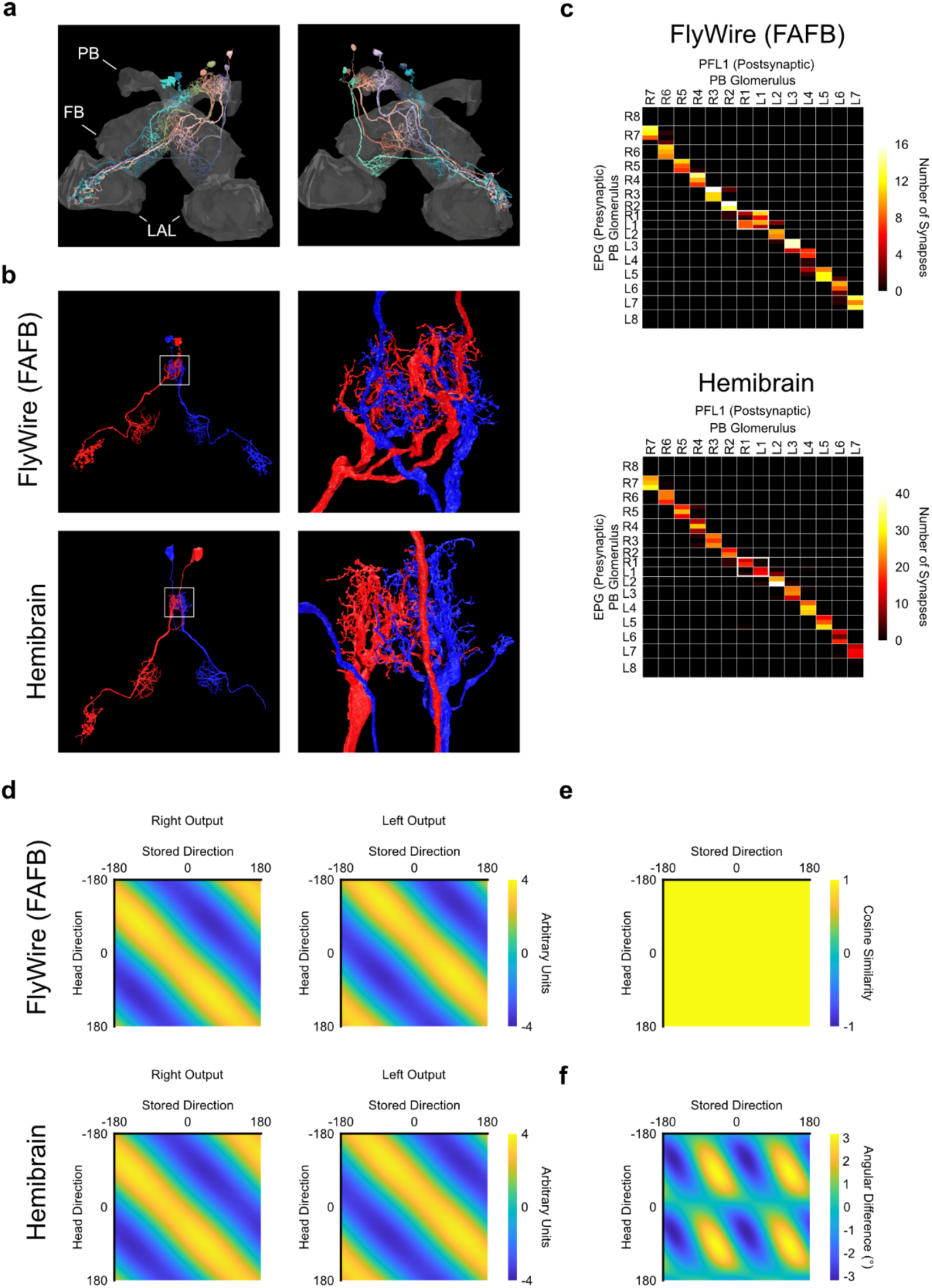
Variation in PFL1 PB tiling. **a,** Rendering of the PFL1 neurons from Flywire. **b,** Renderings of the PFL1 neurons innervating the most medial PB glomeruli from Flywire and the Hemibrain. Notice that in Flywire, both neurons innervate both glomeruli, while in the Hemibrain, each neuron only innervates a single glomerulus. **c,** Connectivity matrix between EPG neurons and PFL1 neurons in Flywire and the Hemibrain. The patterns in PFL1 tiling of the medial two glomeruli are reflected by the connectivity in both datasets (white rectangles at the center). **d,** Output of the circuit model when the preferred head direction assigned to each neuron was based on the tiling pattern from each dataset. **e,** Cosine similarity between the two models in d. **f,** Difference in the direction of the output vectors from the two models in d.

**Extended Data Figure 2:**
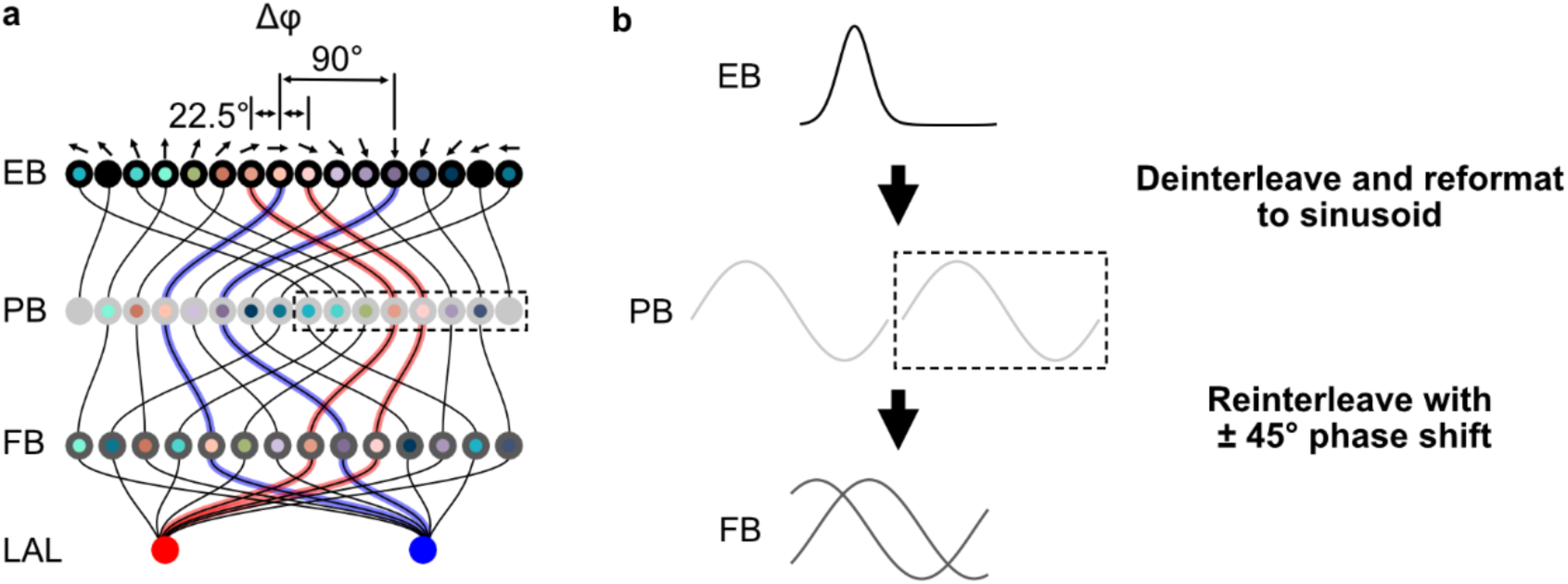
Spatial relationship between the PB and the FB connected by PFL1. **a,** A diagram showing the connectivity of EPG neurons and PFL1 neurons in the central complex. The top row of dots represents wedges in the ellipsoid body. The arrows above the dots represent the preferred head directions of the wedges. Note that neighboring wedges have preferred head directions 22.5° apart. The second row represents glomeruli in the protocerebral bridge. The lines between these rows of dots represent the projection pattern of the EPG neurons from the EB to the PB. The dashed box indicates the left PB. The third row of dots represents the position of the PFL1 dendrites in the FB. The lines connecting the second row to the third row represent the projection patterns of PFL1 dendrites. The bottom row represents the left and right LAL. Right is red; left is blue. The pathways upstream of two neurons from the left and right LALs are highlighted. Notice that in the EB, the two wedges upstream of the two highlighted right neurons flank a neuron with a preferred head direction of 0°. In the FB, however, the same two neurons flank a neuron with a preferred head direction of −90°. **b,** A schematic showing how the EB population code is transformed by each anatomical step.

**Extended Data Figure 3:**
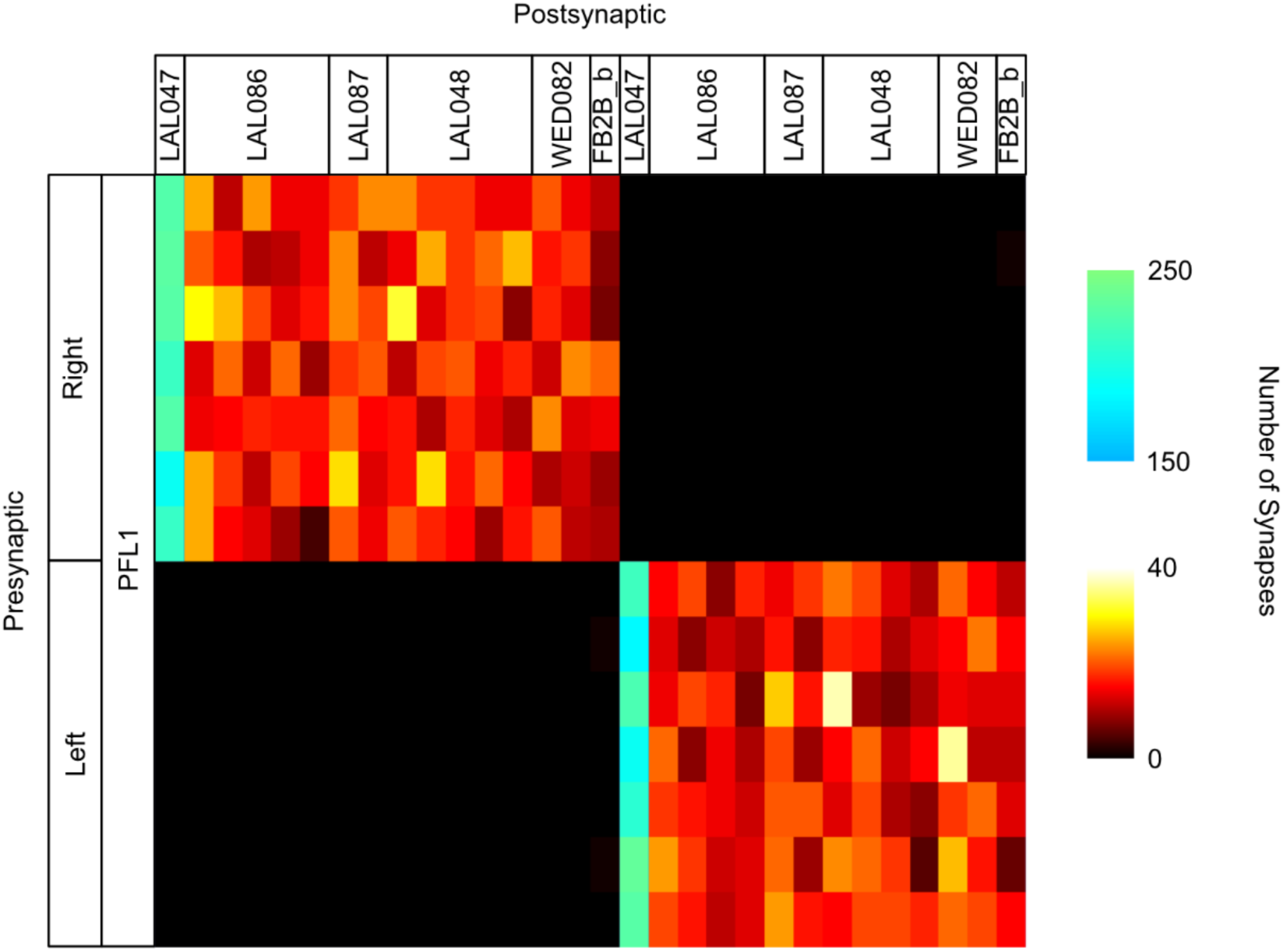
Postsynaptic partners pool input from all PFL1 neurons on that side. Connectivity matrix that shows connections between PFL1 neurons and the top 30 postsynaptic neurons (6 types) by synapse count, sorted by cell type from the Flywire dataset.

**Extended Data Figure 4:**
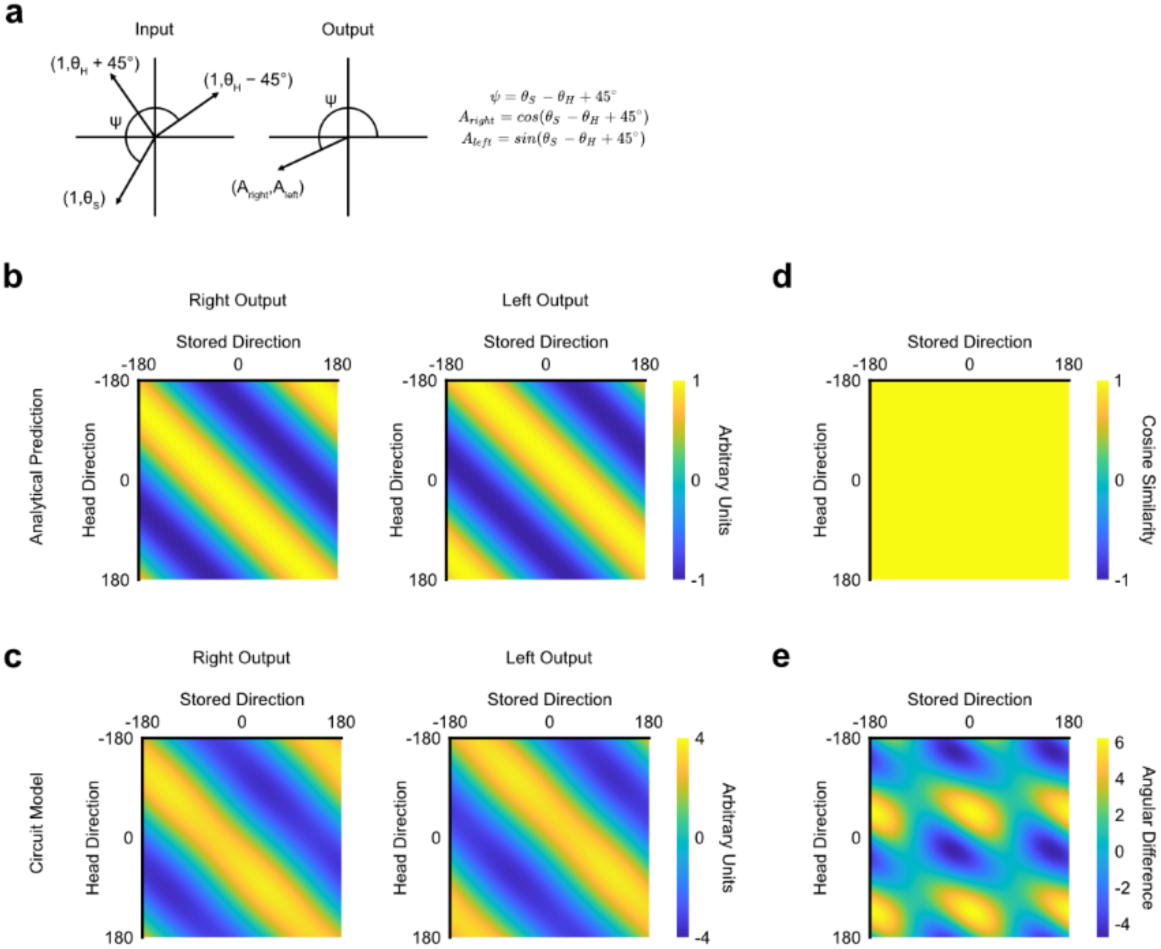
Comparing the circuit model to a true coordinate transformation. **a,** Schematic illustrating how the output vector should be related to the directions of the input vectors if the circuit model does a coordinate transformation. **b,** The x and y coordinates of the output vector from analytically calculating the coordinate transformation. **c,** The output of the circuit model, where the output of each neuron is equal to the product of its inputs. **d,** The cosine similarity between the output of the circuit model in c and the analytical prediction in b. **e,** The difference in direction between the vector defined by the output of the circuit model in c and the analytical prediction in b.

**Extended Data Figure 5:**
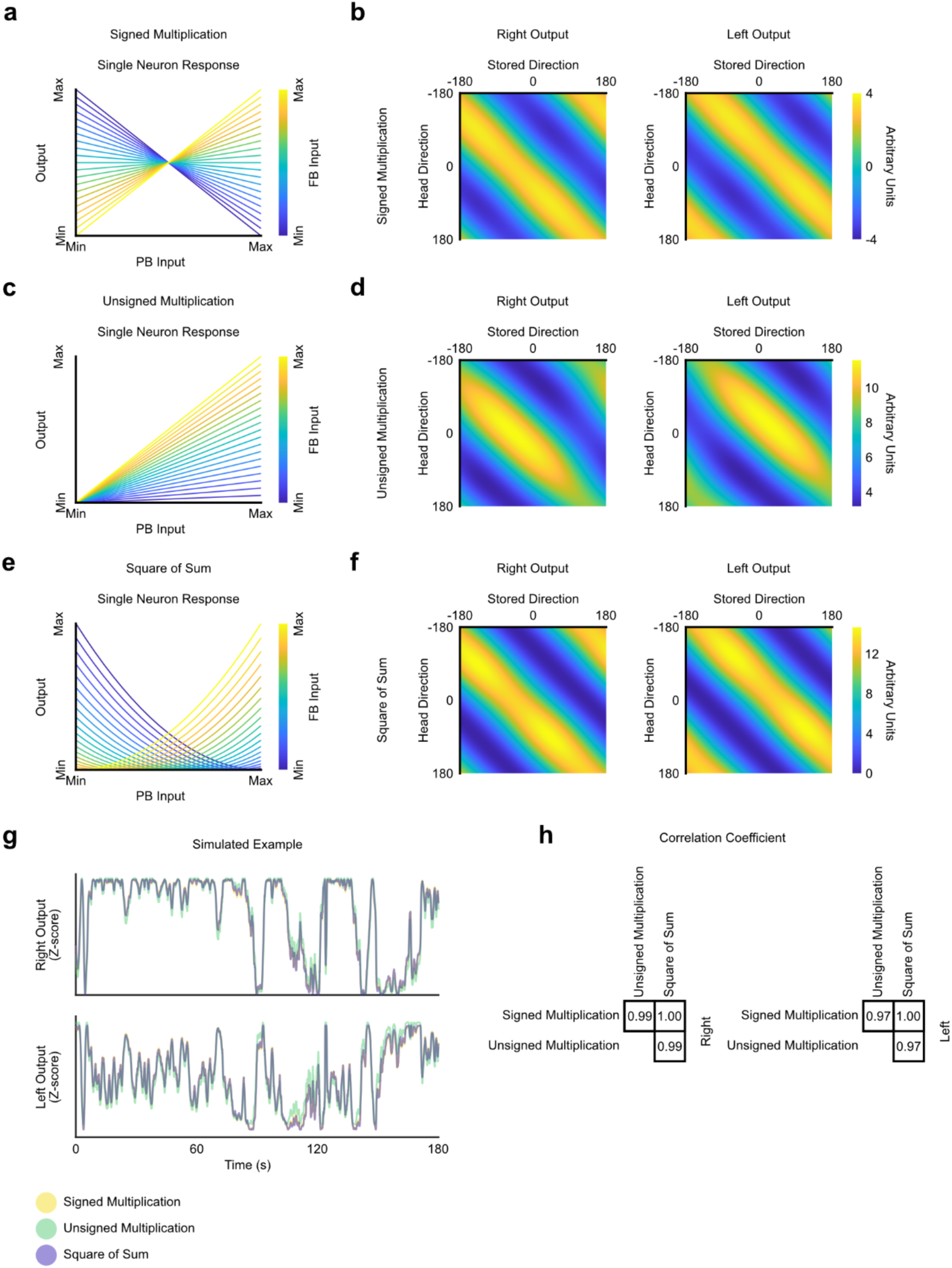
Modeling different activation functions. **a,** Single-neuron response for an activation function that is the product of its input in the PB and the FB. **b,** Output of the circuit model using a multiplicative activation function for the PFL1 neurons. **c,** Single-neuron response for an activation function that is the product of its input in the PB and FB, where the inputs are both positive. **d,** Output of the circuit model using a multiplicative activation function for the PFL1 neurons, where the inputs are positive. **e,** Single-neuron response for an activation function that is the square of the sum of its inputs in the PB and the FB. **f,** Output of the circuit model using the ‘square of the sums’ activation function for the PFL1 neurons. **g,** Simulating the example trial in Fig. 2c using the circuit model with three distinct activation functions that all give the same result. **h,** Correlation coefficients between the simulated example in g.

**Extended Data Figure 6:**
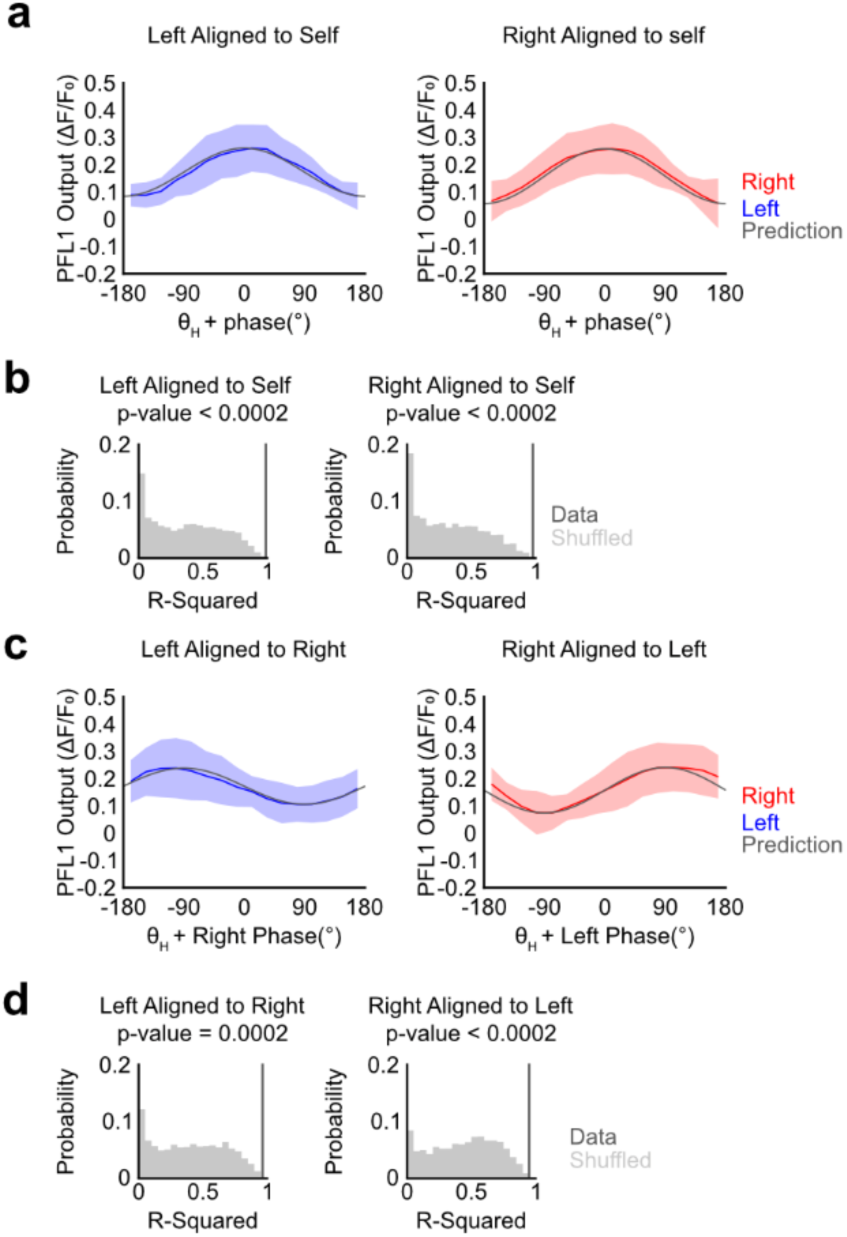
Fitting a cosine function to the PFL1 output. **a,** Comparing PFL1 output to a cosine fit. Half the data for each trial was used to fit a cosine function. The other half of the data is aligned and plotted along with a prediction based on the fitting. Red and blue lines are across-fly averages. The shaded region represents a standard deviation. N=11 flies. **b,** The R-squared value for the data and the predictions in a (dark grey line), compared to a distribution of the same statistic for a shuffled dataset (light grey distribution). **c,** The output of PFL1 aligned to the phase of the opposite side found through fitting a cosine function to the entire trial. Red and blue lines are across-fly averages. The shaded region represents a standard deviation. The grey line is a prediction assuming the two sides are 90° out of phase. N=11 flies. **d,** The R-squared value for the data and the predictions in d (dark grey line) compared to a distribution of the same statistic for a shuffled dataset (light grey distribution).

**Extended Data Figure 7:**
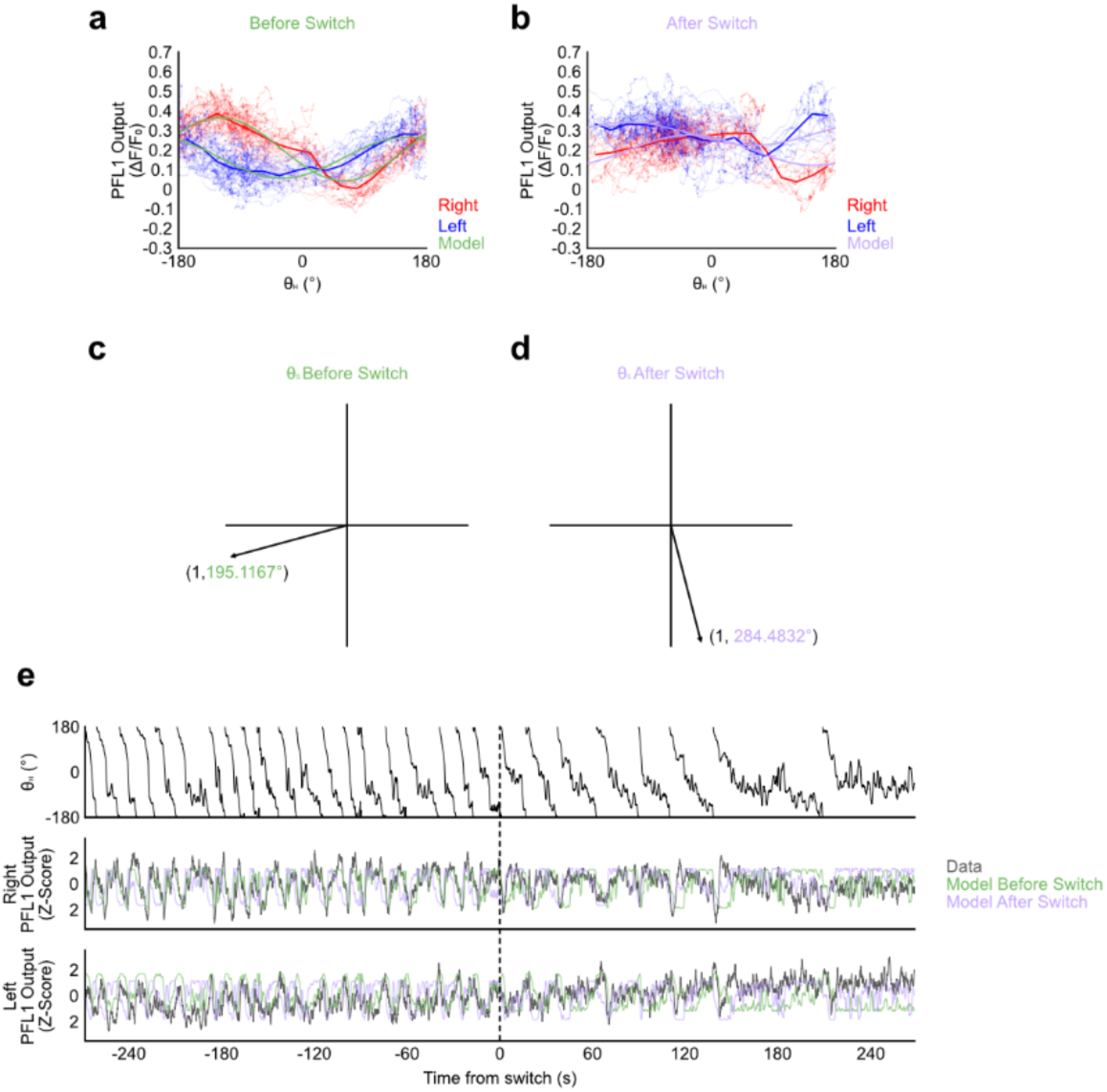
Phase of PFL1 output changes abruptly in a single fly. **a,** PFL1 activity in LALs plotted as a function of the head direction before the change in phase. Green curves are a prediction based on fitting the coordinate transformation model. **b,** PFL1 activity plotted as a function of the head direction after the change in phase. Purple curves are a prediction based on fitting the model. **c,** The stored direction found by fitting the model to the data prior to the switch. **d,** The stored direction found by fitting the model to the data after the switch. **e,** Time series centered on the moment the phase switches. Top: head direction. Middle: Right LAL recording. Bottom: Left LAL recording. The green and purple traces are simulations from the circuit model based on the head direction for each time point and the stored directions from before and after the switch, shown in c and d.

**Extended Data Figure 8:**
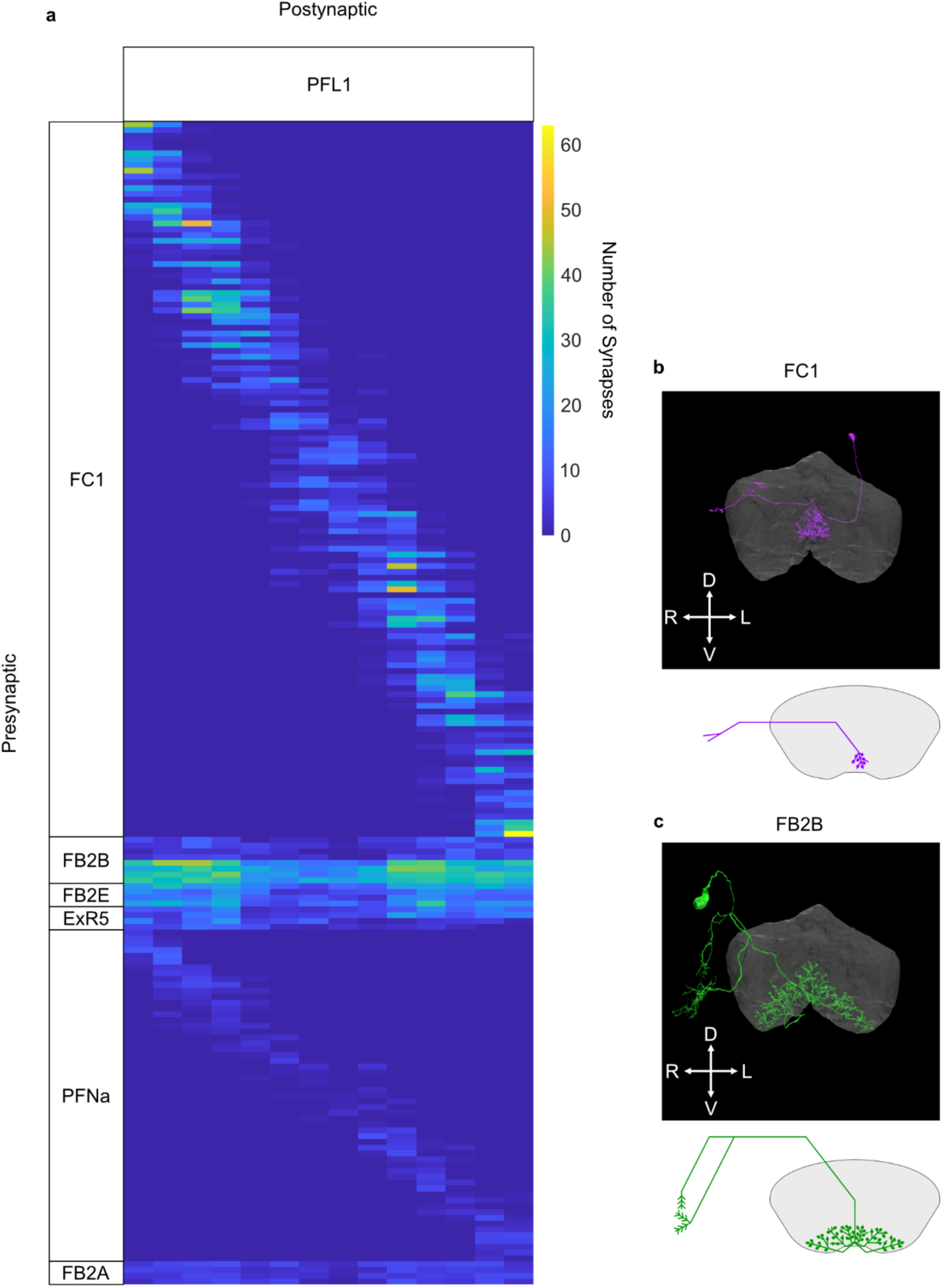
FB Input to PFL1. **a,** A connectivity matrix showing the number of synapses in the FB between presynaptic neurons from the top presynaptic cell types and PFL1 in the hemibrain dataset. Note that neurons belonging to these cell types that have no synapses with PFL1 in the FB are excluded from this plot. **b,** The image of an individual FC1 neuron from the hemibrain dataset (top). Drawing of an individual FC1 neuron (bottom). Note that FC1 neurons are columnar neurons, so they occupy a narrow portion of the FB. **c,** The same as b, but for an individual FB2B neuron. Note that this neuron is a tangential neuron, so it occupies the full width of the FB. D, dorsal. L, left. V, ventral. R, right.

**Extended Data Figure 9:**
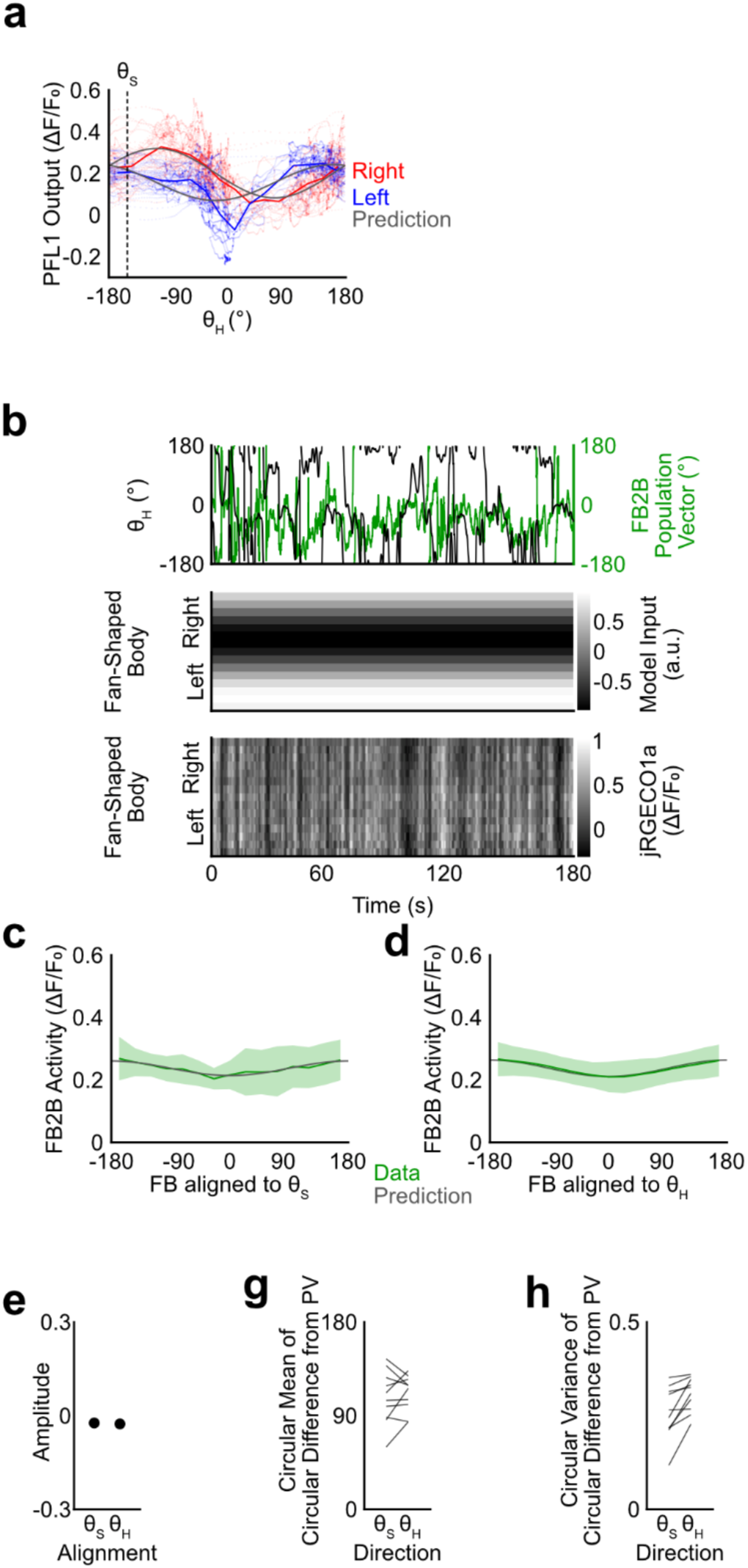
FB2B calcium activity is uniform across the FB. **a,** Data from an example trial, showing the output of PFL1 plotted as a function of the head direction, along with a prediction from fitting the coordinate transformation model to the data. Dots are time points. Red and blue lines are the averages of the dots. The dotted line indicates the stored direction found through fitting. **b,** Data from the example trial in a. Top: the head direction and the population vector of FB2B. Middle: The simulated FB input from the coordinate transformation model, predicted based on fitting. Bottom: Calcium recording from FB2B in the FB. **c,** FB2B activity plotted as a function of the position in the fan-shaped body aligned to the stored direction. The green line reflects the across-fly mean. The shaded region represents a standard deviation. The dark grey line shows the predicted result if the phase were to encode the stored direction with an amplitude fit to the data. N = 10 flies. **d,** The same as c except aligned to the head direction. **e,** The amplitude found by fitting the prediction. **f,** The circular mean of the circular difference between the FB2B population vector and the head direction or the stored direction. **g,** The circular variance of the circular difference between the FB2B population vector and the head direction or the stored direction.

**Extended Data Figure 10:**
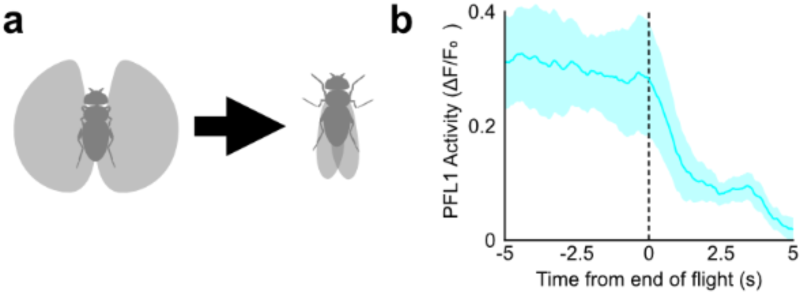
PFL1 activity is flight-dependent. **a,** A schematic illustrating the transition from flying to not flying. **b,** Calcium activity in the axons of PFL1 when the fly stops flying. The shaded region represents a 95% confidence interval. N=26 LAL recordings from 13 flies.

**Supplementary Video 1:** Anatomy of PFL1

The video starts with a 360° rotation. Then, the video focuses on the fan-shaped body dendrites. Neurons with axons on opposite sides of the brain are pulled apart to reveal that each population tiles the fan-shaped body independently. Next, we bring a single neuron with axons on the left to the center to reveal the neurons from the right with which it overlaps. Neurons appear one by one, showing that the dendrites of each neuron are sandwiched between two neurons from the opposite side of the brain.

## Mathematical Appendix

We show the similarity of three models: the ‘unsigned multiplication’ model, the ‘square of sum’ model, and the ‘signed multiplication’ model.

### Unsigned Multiplication

The activation function for the unsigned multiplication model is:

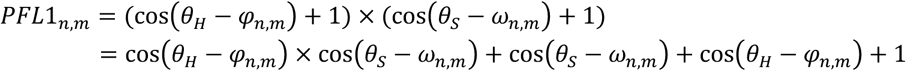

The output of the model sums the activity of all the neurons on each side of the brain:

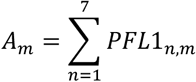

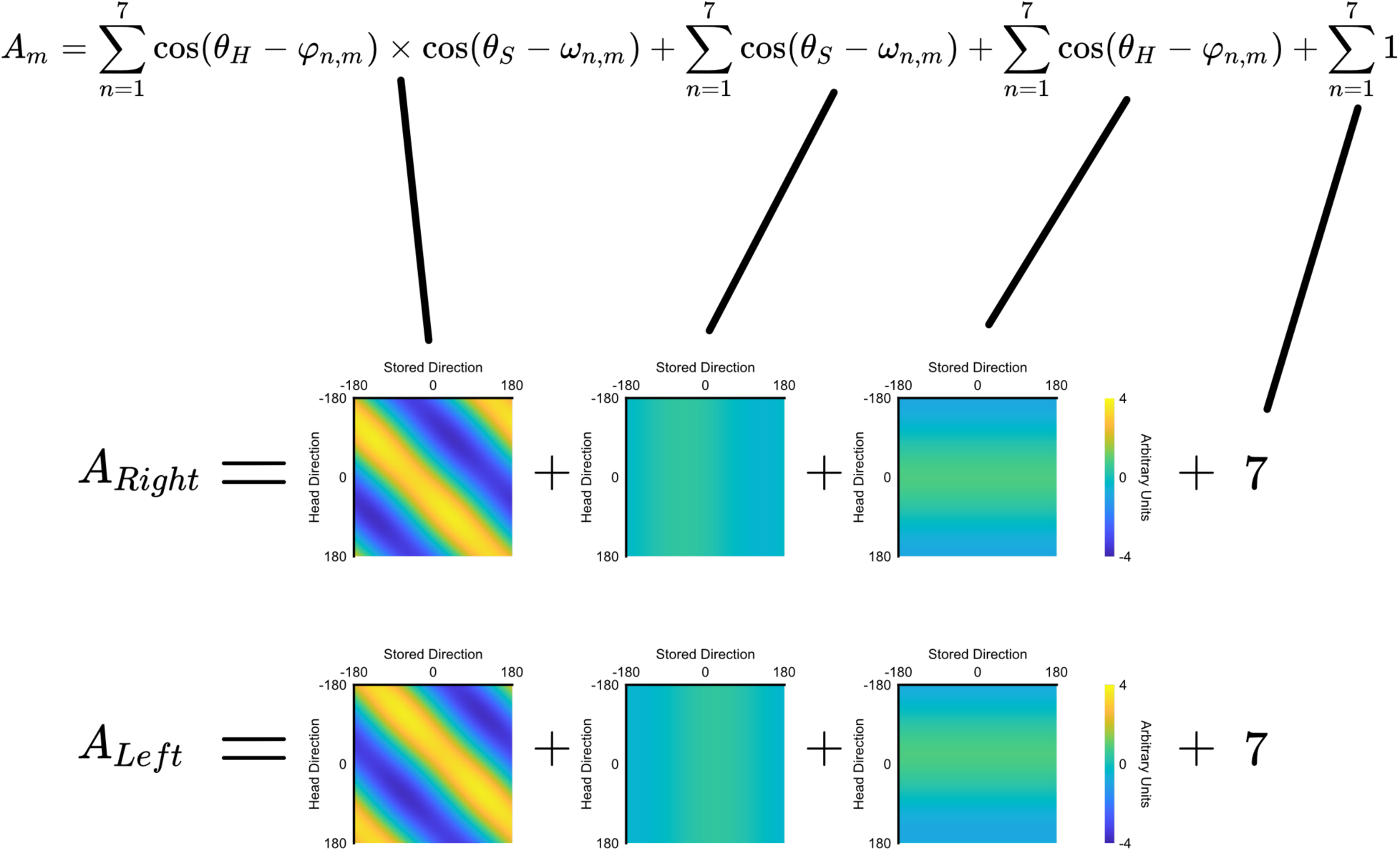

The second and third terms are close to zero, and thus the result is nearly identical to the signed multiplication model with a bias.

### Square of Sum

The activation function for the square of sum model:

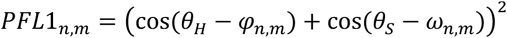

Using trigonometric identities, we can rewrite the equation:

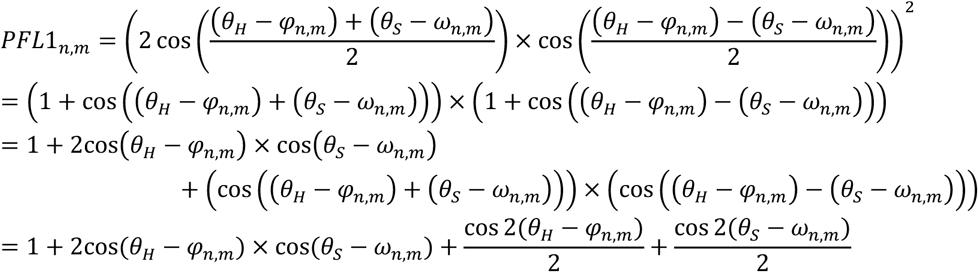

The output of the model sums the activity of all the neurons on each side of the brain:

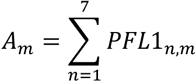

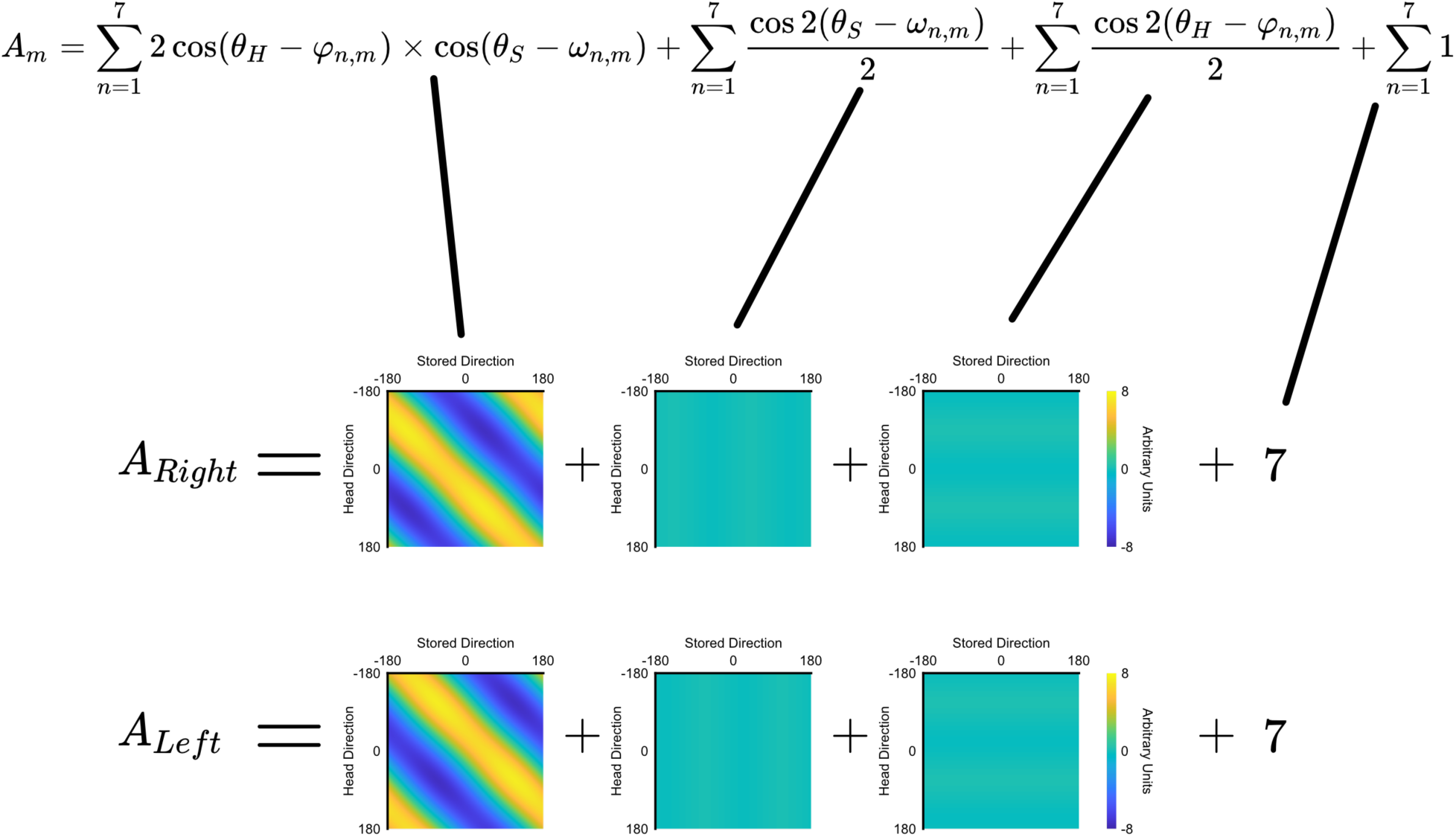

Again, the second and third terms are close to zero. The output is nearly identical to the signed multiplication model except with scaling (x2) and a bias.

The second and third terms are small in both models, because the preferred head and stored directions are spread out across a period, which makes the sum of these terms largely cancel out. If the preferred directions were evenly spaced across the period, the sum would equal exactly zero.

We used Pearson’s correlation as a measure of similarity between the data and the model, which is not affected by linear scaling and biasing. Therefore, all three models produce nearly identical results.

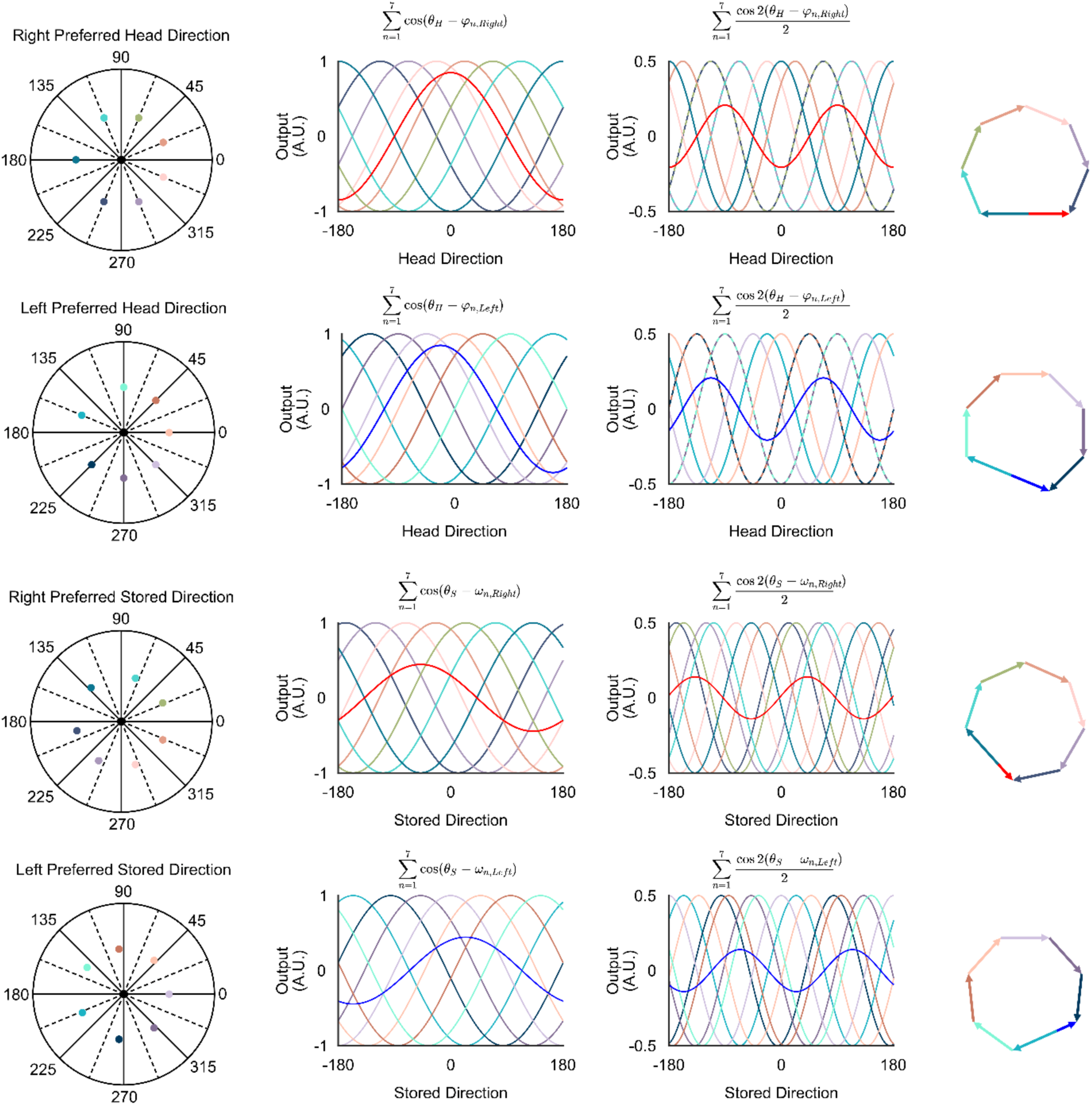

Red and blue curves or arrows represent the sum of each term across neurons. Other colored curves and arrows represent the value of the term for individual neurons.

## Notes

### Competing Interest Statement

The authors have declared no competing interest.

